# Structural analysis of the NifL-NifA complex reveals the molecular basis of anti-activation of nitrogen fixation gene expression in *Azotobacter vinelandii*

**DOI:** 10.1101/2025.06.05.658055

**Authors:** Marcelo Bueno Batista, Jake Richardson, Michael W. Webster, Dmitry Ghilarov, John W. Peters, David M. Lawson, Ray Dixon

## Abstract

Understanding the molecular basis of regulated nitrogen (N_2_) fixation is essential for engineering N_2_-fixing bacteria that fulfill the demand of crop plants for fixed nitrogen, reducing our reliance on synthetic nitrogen fertilizers. In *Azotobacter vinelandii* and many other members of Proteobacteria, the two-component system NifL-NifA controls the expression of *nif* genes that encode the nitrogen fixation machinery. The NifL-NifA system evolved the ability to integrate several environmental cues, such as oxygen, nitrogen, and carbon availability. The nitrogen fixation machinery is thereby only activated under strictly favorable conditions, enabling diazotrophs to thrive in competitive environments. Whilst genetic and biochemical studies have enlightened our understanding of how NifL represses NifA, the molecular basis of NifA sequestration by NifL depends on structural information on their interaction. Here, we present mechanistic insights into how nitrogen fixation is regulated by combining biochemical and genetic approaches with a low-resolution cryo-EM map of the oxidized NifL-NifA complex. Our findings define the interaction surface between NifL and NifA and reveal how this interaction can be manipulated to generate bacterial strains with increased nitrogen fixation rates able to secrete surplus nitrogen outside the cell, a crucial step in engineering improved nitrogen delivery to crop plants.

## Introduction

To meet current demands for global food production, agriculture relies on synthetic nitrogen fertilizers. Nitrogen fertilizer enabled the Green Revolution and has been one of the main drivers for exponential population growth since the mid-50s. However, synthetic nitrogen fertilizer production and distribution accounts for a large proportion of energy use worldwide, consuming 3%–5% of the world’s natural gas production [1–5]. Currently, an estimated 12 Gt/year of CO_2_ equivalents is released into the atmosphere from the industrial production of ammonia. This represents around 1.5% of all greenhouse gas emissions globally [4, 6]. The engineering of more efficient delivery of fixed nitrogen by free-living soil bacteria that can associate with cereal crops [7, 8] is a viable alternative to current practices, enabling more sustainable agriculture. In *Azotobacter vinelandii* and many other Proteobacteria, the expression of *nif* genes encoding the machinery for biological nitrogen fixation is controlled by the non-canonical two-component system NifL-NifA [8–10]. In the NifL-NifA system, the regulatory signal transduction does not involve phospho-transfer between the two components, as NifL does not have phosphorylation activity, and NifA does not have a receiver domain [11]. Instead, regulatory outputs are driven by stoichiometric interactions between NifL, NifA, and the global signal transduction protein, GlnK, in response to the oxygen, nitrogen, and carbon status in the cell [8–10].

NifA is a σ^54^-dependent bacterial enhancer binding protein (bEBP) that activates transcription of the nitrogen fixation (*nif*) genes [12]. Its central domain comprises the AAA+ ATPase domain responsible for ATP hydrolysis-dependent stimulation of open-complex formation at σ^54^-dependent promoters. NifA also has a HTH DNA-binding domain at its C-terminus, and a regulatory GAF domain at its N-terminus, which can bind 2-oxoglutarate to mediate the carbon signal transduction response. NifL is an anti-activator protein homologous to sensor histidine kinases in canonical two-component systems that does not hydrolyse ATP or function as a kinase [11]. In its N-terminus, NifL comprises two in tandem PAS domains, namely PAS1 and PAS2, whilst the C-terminus contains kinase-like DH and GHKL domains [9, 13]. Under oxygen-excess conditions, the FAD moiety within the PAS1 domain of NifL is oxidized [14, 15], leading to a series of conformational changes that allow NifL-NifA complex formation and thereby inhibits promoter activation by NifA by preventing its oligomerization (Fig. 1). NifL has a second PAS2 domain that does not bind FAD and instead is involved in transmitting conformational changes from the PAS1 domain to the DH and GHKL domains in the NifL C-terminus [16–18]. The DH domain, referred to as the DHp (dimerization and histidine phosphotransferase domain[13]), in histidine protein kinases involved in NifL oligomerization, whilst the GHKL domain regulates NifL activity by binding either ADP and ATP [19, 20]. As the K_d_ for ADP is at least 10x higher than that of ATP, NifL acts as a sensor of the energetic status of the cell in which binding of ADP stimulates the interaction with NifA under conditions unfavorable for nitrogen fixation.

**Figure 1.**
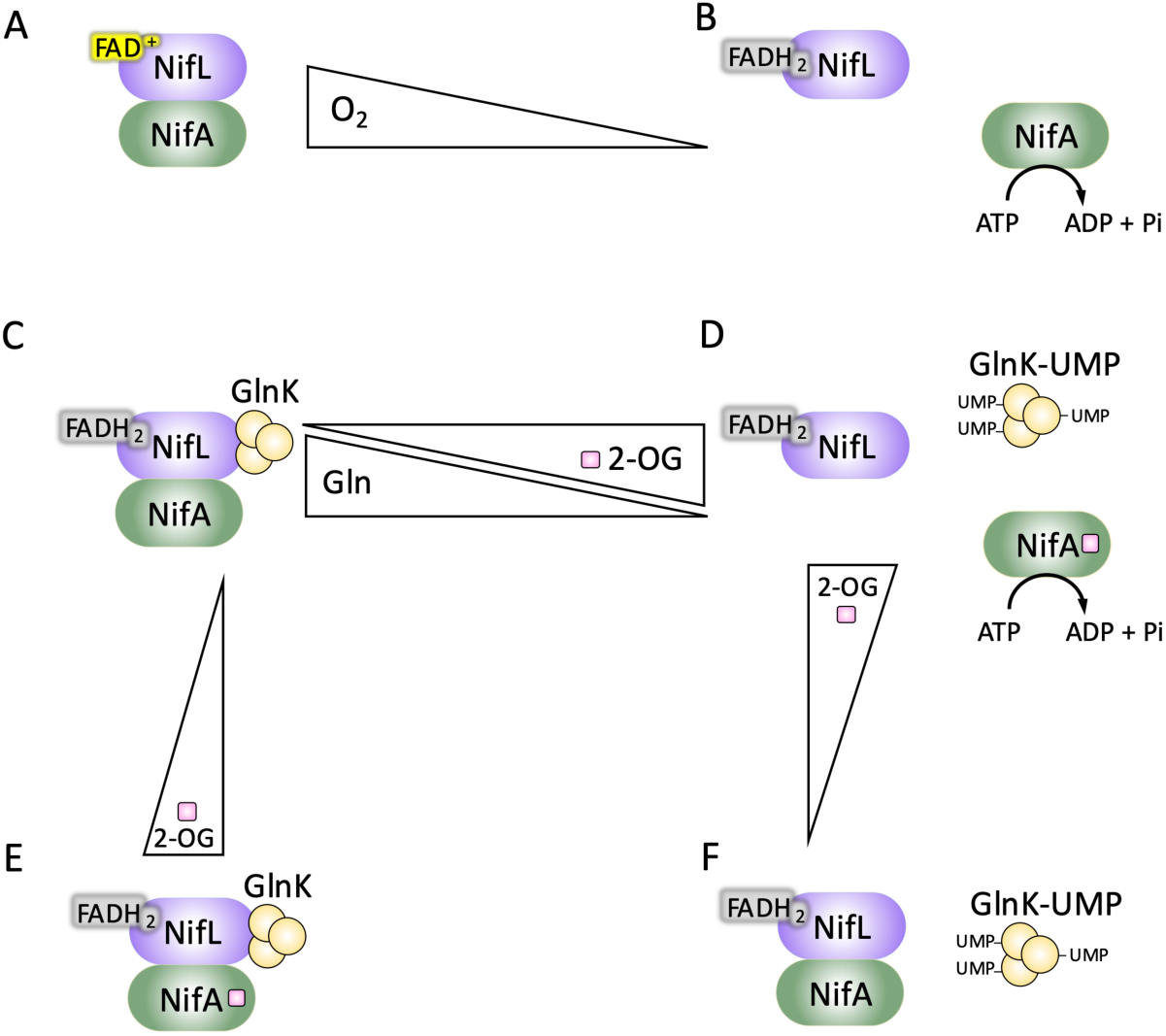
Current model for NifL control of NifA activity dependent on different environmental signals in *A. vinelandii*. (A) Under excess oxygen conditions, the FAD cofactor in the NifL PAS1 domain is oxidized, enabling NifL to adopt an inhibitory conformation that is competent to inactivate NifA. (B) Under limiting oxygen conditions, the FAD cofactor is fully reduced (FADH2), triggering conformational changes in NifL that prevent interaction with NifA, releasing its activity. In the reduced state, NifL can still inhibit NifA in response to the levels of nitrogen and carbon as signaled by glutamine (Gln) and 2-oxoglutarate (2-OG), respectively. (C) Under nitrogen excess conditions, non-uridylylated GlnK stimulates the formation of the GlnK-NifL-NifA complex that inhibits NifA activity. (D) Upon a shift to low nitrogen (signaled by low Gln levels), GlnK is uridylylated (GlnK-UMP). Under optimal carbon supply, a reduction in Gln is accompanied by an increase in 2-OG levels. Under these conditions, the NifA GAF domain is saturated with 2-OG, allowing for the complete dissociation of the complex and the release of NifA activity. (E) Under nitrogen excess conditions, the GlnK-NifL-NifA ternary complex is stable even if the GAF domain in NifA is saturated with 2-OG in carbon-rich conditions. (F) Under nitrogen limitation, if 2-OG is limiting, an inhibitory binary complex between NifL and NifA can still be formed.

Regulation of NifA activity by nitrogen and carbon levels in organisms that encode NifL also requires the PII signal transduction protein GlnK (Fig. 1). In *A. vinelandii,* under conditions of nitrogen excess, a ternary GlnK-NifL-NifA complex is formed to inhibit NifA activity depending on the GlnK uridylylation state relative to the nitrogen status of the cell (for comprehensive reviews, see [21–24]). A NifL-NifA complex can still be formed when NifL is reduced and GlnK is uridylylated under nitrogen-limiting conditions if the cellular carbon levels, and hence the 2-oxoglutarate levels, are low. The binding of 2-oxoglutarate, an allosteric effector at the interface of carbon and nitrogen metabolism [25], to the GAF domain of NifA is required for the activator to escape inhibition by NifL [8, 10, 26] (Fig. 1). This provides an additional layer of complexity to the NifL-NifA regulation that ensures activation of nitrogen fixation, a highly energy demanding process, is only triggered under carbon-rich conditions; for example, when the bacterium closely interacts with crop plant roots and can benefit from carbon-rich exudates.

Biochemical and genetic studies have provided important insights into how the NifL-NifA system integrates diverse regulatory inputs to regulate nitrogen fixation [8–10, 26]. However, efforts to manipulate this system predictably to engineer diazotrophs with enhanced ability to fix nitrogen and release ammonia to benefit crop nutrition have been hampered by the lack of structural information. We recently obtained structural snapshots of the dynamic structural changes in NifL that lead to the inhibition of NifA using SEC-SAXS and protein labeling followed by mass spectrometry [27]. Although methods of NifL-NifA complex purification have been known for decades [28, 29], structural characterization of this complex has remained elusive.

In this study, we obtained a low-resolution cryo-EM reconstruction of the NifL-NifA complex that is consistent with a confident structural prediction. Extensive mutagenesis allowed us to confirm the interaction interface between NifL and NifA that provides insight into the inhibitory conformation of NifL that can hold NifA in an inactive state. This insight allowed reinterpretation of previously obtained mutant phenotypes and enabled the isolation of novel variants with sought-after properties in *A. vinelandii*. Substitutions of amino acids comprising the surface of interaction in either NifL or NifA led to de-regulated expression of nitrogenase, resulting in the release of excess ammonia outside the bacterial cell. This underpins a strategy to engineer synthetic associations between associative diazotrophs and plants that deliver fixed nitrogen directly to cereal crops.

## Results

### Purification and structural characterization of the NifL-NifA complex

Following our previous SEC-SAXS-guided analysis of the structural dynamics of *A. vinelandii* NifL dimer in the oxidized and reduced states bound either to ADP or ATP [27], we sought to determine the structure of the oxidized ADP-bound NifL dimer in complex with NifA to get further insights into how NifL represses activation of nitrogen fixation genes. A previously established protocol for NifL-NifA preparation [28, 29] was optimised to achieve high yields of the protein complex under oxidizing conditions and in the presence of ADP. Affinity chromatography and two rounds of size exclusion chromatography (SEC) were found to improve protein complex homogeneity (Fig. 2).

**Figure 2.**
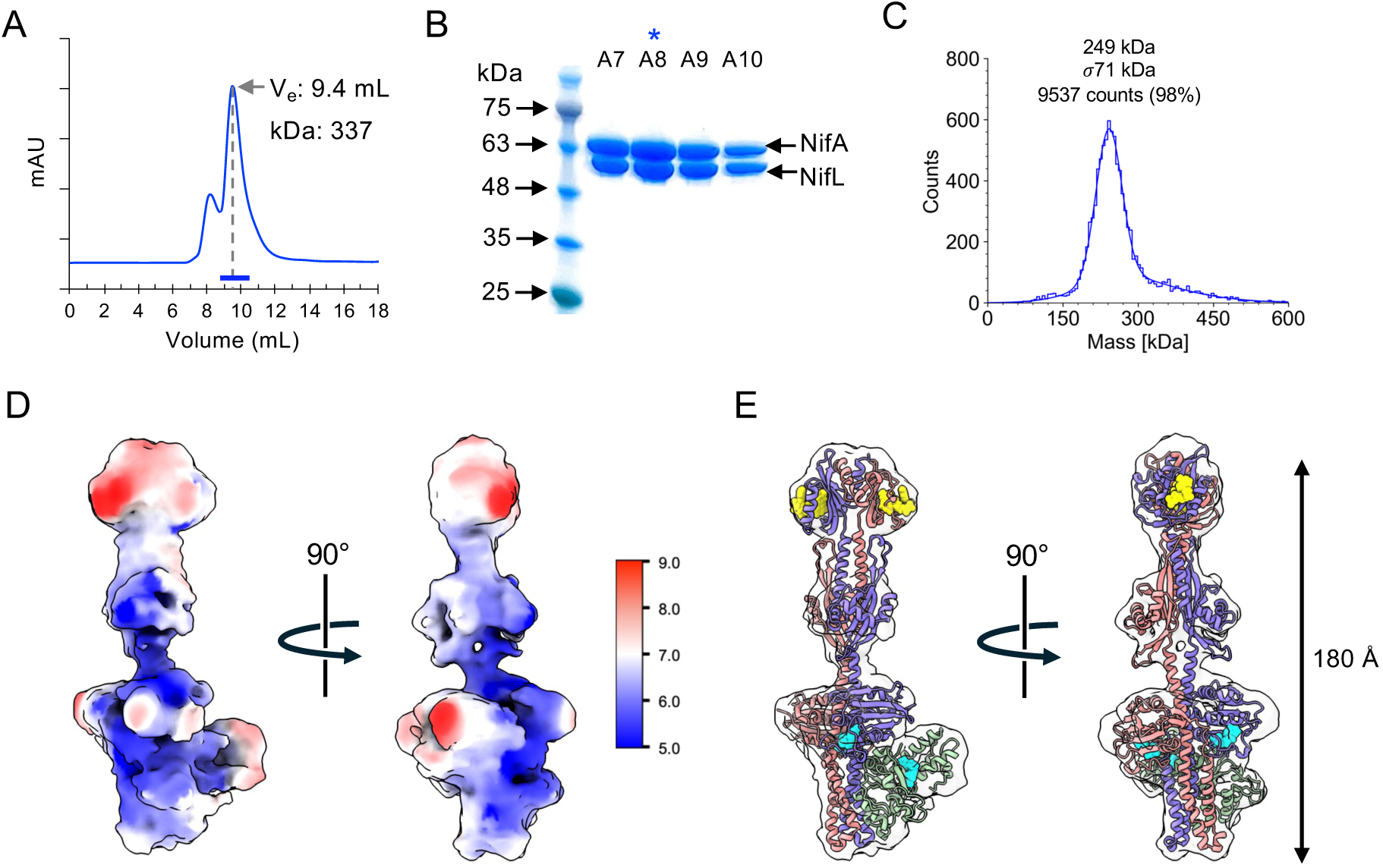
Purification and structural analysis of the oxidised NifL-NifA complex, prepared in the presence of ADP. (A) size exclusion chromatography (SEC) elution profile of the NifLA complex. The thick blue line in the X-axis indicates the fractions loaded for SDS-PAGE visualisation. (B) SDS-PAGE from SEC elution fractions in A. The blue asterisk indicates the fraction used for mass photometry analysis. (C) Mass photometry analysis of the purified NifLA complex. (D) Orthogonal views of the cryo-EM reconstruction having a global resolution of 6.45 Å with the local resolution plotted onto the surface. (E) Cryo-EM reconstruction displayed as a semi-transparent surface together with the refined model comprised of two copies of NifL (salmon and purple cartoons) and a single NifA AAA+ domain (green cartoons). Also shown as cyan van der Waals spheres are the bound cofactors and nucleotides. See Figure 2 for more detail.

The peak fraction of the SEC eluate was analysed by mass photometry to assess NifL-NifA complex stoichiometry (Fig. 2C). The main observed mass distribution was approximately 240 kDa, consistent with a NifL-NifA complex comprising two monomers of NifL and two monomers of NifA. This mass estimate contrasts with a previous assessment suggesting that the complex was formed by a tetramer of NifL and a tetramer of NifA [28]. As discussed previously, for the structural characterization of *A. vinelandii* NifL [27], we propose that analytical SEC overestimates the molecular mass due to the elongated complex structure described below.

We sought to determine the structural architecture of the purified NifL-NifA complex by single-particle cryo-EM. Two-dimensional class averages revealed an elongated particle ∼180 Å in length, which is comparable to the previous structural characterization of the NifL dimer alone (Fig. S1) [27]. After further processing, a reconstruction was obtained to an average resolution of ∼6.5 Å (Fig. S1, Fig. 2D, E). However, the generation of a high-resolution 3D volume for the complex was beyond the limitations of the data, since the particle images were noisy, displayed conformational heterogeneity, and suffered from orientational bias. The reconstruction was consistent with part of a structural prediction of NifL-NifA complex generated with 2:2 stoichiometry using AlphaFold3 (Fig. S2). In the structural prediction, two NifL subunits are in a two-fold symmetric arrangement. The extended central axis is comprised predominantly of intertwined α-helices with a pair of PAS1 domains at one end of the complex, a pair of PAS2 domains at the midpoint, and a pair of GHKL domains at the opposing end (Fig. 3). The reconstruction revealed significantly less density for NifA, and we were only able to model the AAA+ domain of a single subunit into the density, interacting with the GHKL and the central DH domains of NifL. This domain is predicted to be the only part of NifA that interacts directly with NifL in the model. It has the highest pLDDT scores, and the PAE plot shows reasonable confidence in the placement of this domain with respect to the C-terminal portion of NifL. Whilst the N-terminal GAF domain and the C-terminal HTH domain of NifA are predicted to be structured, the PAE plot indicates no confidence in their placement with respect to the AAA+ domain or the rest of the complex. However, the lack of a resolvable second NifA subunit is intriguing. The AlphaFold3 model predicts that the N-terminal GAF domain and C-terminal HTH domain of NifA are not anchored to the core of the complex. Could one or both of these apparently mobile domains have a tendency to associate with the air-water interface and either initiate the complete unfolding of the NifA subunit or simply detach it from the complex. Given that the two NifA subunits lie on opposing sides of the complex, perhaps in the majority of cases, only one NifA subunit per particle is affected, thereby giving rise to the asymmetric particles we observed.

**Figure 3.**
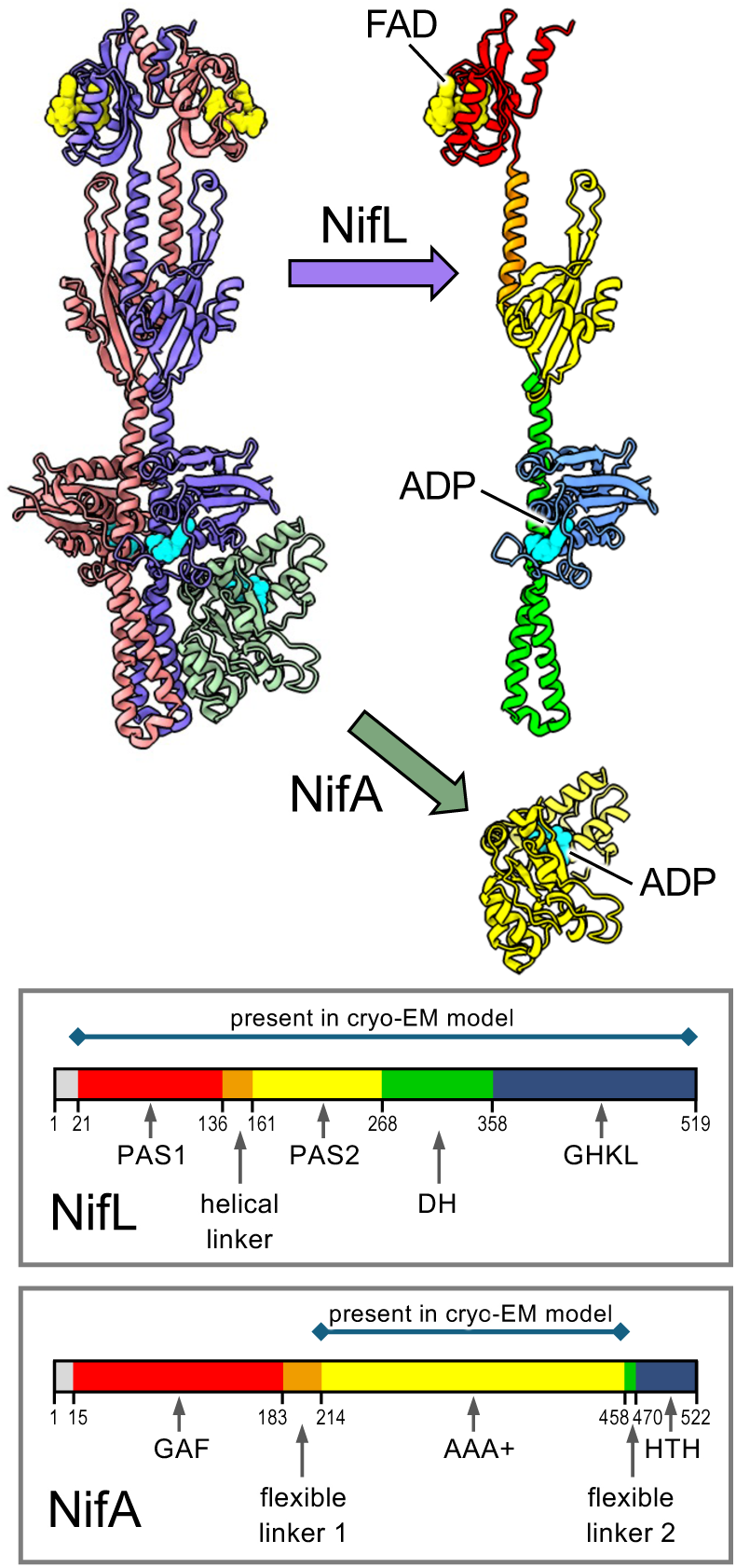
Detail of the refined NifL-NifA cryo-EM structure. Two copies of the NifL sequence could be fitted to the density (salmon and purple cartoons), with the exception of the first twenty residues and the last two, forming a two-fold symmetric arrangement. Only a single copy of the NifA AAA+ domain (residues 210-458; green cartoons) could be modelled interacting simultaneously with the NifL GHKL domain and the DH domain. The separated subunits on the right are coloured by domain in accordance with the respective schemes in the panels below. The locations of the FAD and ADP ligands are indicated (yellow and cyan van der Waals spheres respectively). These were predicted from the starting AlphaFold3 model and assumed to be present, but there was insufficient detail in the 6.45 Å-resolution reconstruction for them to be resolvable in the density. Note the distinctive constriction roughly at the junction of the upper and lower halves formed by opposing helices from the DH domains of the two NifL subunits; this is clearly visible in the 2D classes (Fig. S1).

Despite the limited resolution of our cryo-EM model analysis, it nevertheless provides orthogonal, experimental support for the predictive models generated by AlphaFold3 and, therefore, a robust three-dimensional framework for the interpretation of the wealth of genetic and biochemical data on this system. The overall architecture of the NifL-NifA complex provides a structural explanation for the sequestration and inhibition of NifA by NifL. The observed interaction with NifL precludes the formation of the predicted NifA hexamer that is required to activate the transcription of nitrogen fixation genes. AlphaFold models suggested that the architecture of the NifL-NifA complex is conserved across the phylogenetic spectrum of nitrogen-fixing proteobacteria that utilize this regulatory system, enabling the identification of conserved amino acid motifs in the interaction surface between NifL and NifA pairs in diverse diazotrophs (Fig. S3).

### Biochemical and genetic characterization of the NifL-NifA interaction surface

To confirm that the DH and GHKL domains of NifL and the AAA+ domain of NifA are sufficient for the NifL-NifA interaction, a truncated NifL_269-519_-NifA_188-445_ complex was purified to homogeneity in the presence of ADP (Fig. 4A). In contrast to observations with the full-length complex, the estimated mass inferred from SEC correlated well with mass photometry (approximately 120 kDa), indicative of a more compact, globular structure of the truncated components as predicted from the structural model. Consistent with the structural model, full-length NifA co-purified with truncated NifL lacking the N-terminal PAS domains (NifL_269-519_) (Fig. 4B), and full-length NifL co-purified with truncated NifA containing only the AAA+ domain (NifA_188-445_) (Fig. 4C).

**Figure 4.**
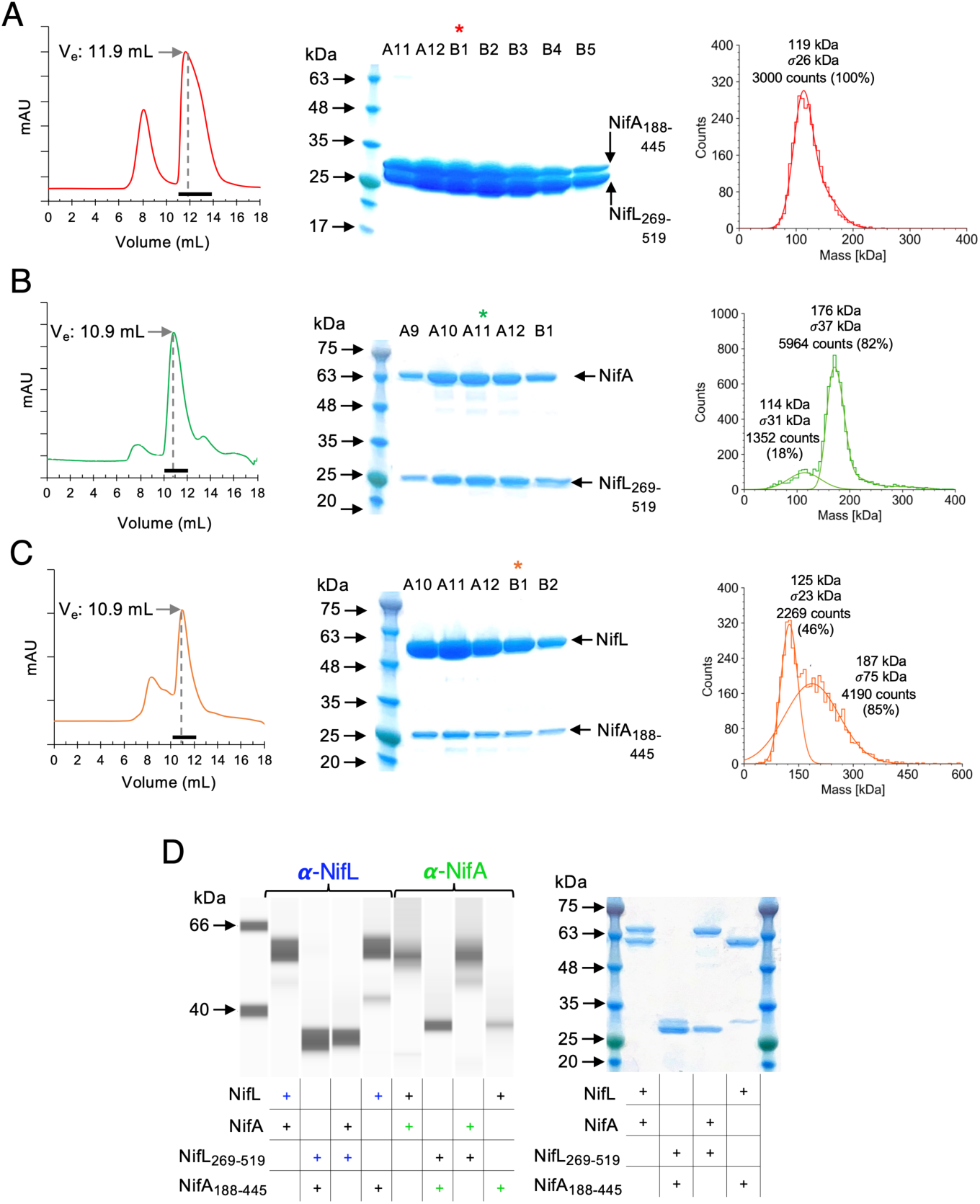
Purification of truncated versions of the NifL-NifA complex allows identification of the surfaces essential for the interaction. (A) Co-purification of NifL269-519 and NifA188-445. (B) Co-purification of NifL269-519 and NifA. (C) Co-purification of NifL and NifA188-445. (A-C) Left, size exclusion chromatography (SEC) elution profile of the purified complexes. The thick black line in the X-axis indicates the fractions loaded for SDS-PAGE visualisation. Middle, SDS-PAGE from SEC elution fractions in the left. The asterisks indicate the fractions used for mass photometry analysis. Right, mass photometry analysis of the purified complexes. Size exclusion chromatography profile of various purified NifL-NifA complexes compared to the molecular weight calibration standards is available in Fig. S4. (D) Immunodetection of purified complexes using polyclonal antisera raised against Av-NifL and Av-NifA. Left, automated western blot image. 0.2 ug of protein was loaded per well/capillary. Right, mirror SDS-PAGE analysis of samples used for immunodetection. 4 ug of protein was loaded per lane.

To understand the functional contribution of individual residues at the predicted NifL-NifA interface, we performed extensive site-directed mutagenesis guided by the structural model. Selection of residues for substitution was further guided by the identification of conserved amino acid motifs in the predicted interaction surface between NifL and NifA pairs in diverse diazotrophs, since AlphaFold models suggested the architecture of the NifL-NifA complex is likely conserved across the phylogenetic spectrum of nitrogen-fixing proteobacteria that use this regulatory system (Fig. S3). An overview of the *A. vinelandii* NifL-NifA interface is presented in Fig. 5A with substituted amino acids indicated. Residues close to the nucleotide-binding pockets of each subunit were avoided as they likely impart functional roles other than only complex formation (Fig. 5B-C) [12, 30].

**Figure 5.**
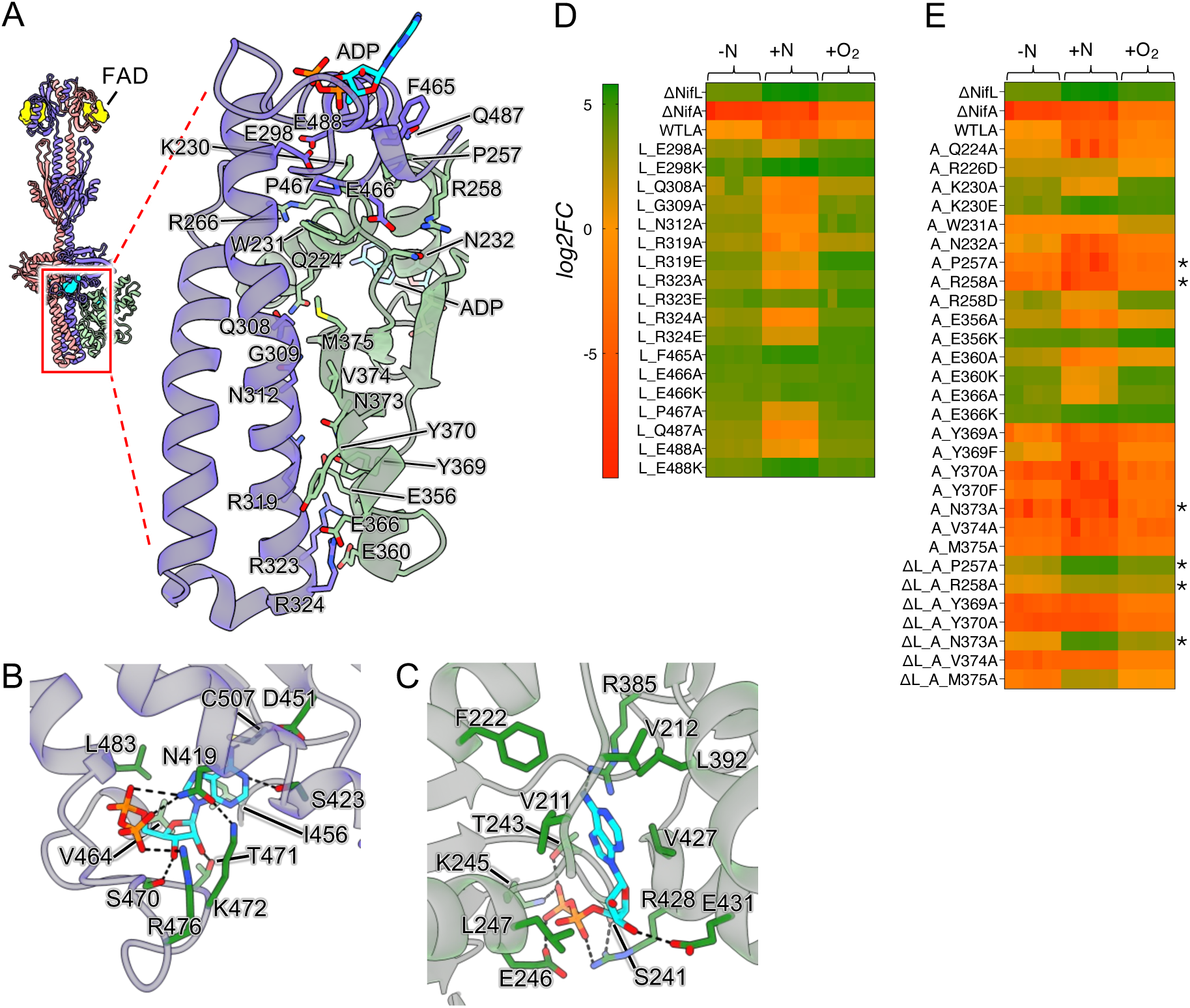
Analysis of residues important for the interaction between NifL and NifA. (A) Close-up of the region targeted for mutagenesis with every substituted residue shown in stick representation and labelled. For clarity, extraneous structure in the foreground and background has been omitted and the backbone cartoons are shown semi-transparent (NifL, purple; NifA, green). (B and C) Predicted nucleotide binding sites for NifL and NifA respectively. Importantly, the key residues shown do not overlap with those that were substituted. (D and E) β-galactosidase activity of wild-type and mutant NifL and NifA variants, respectively, measured in *E. coli* ET8000 carrying a *PnifH::lacZ* reporter. Activity is presented as log2FC in relation to the activity of the wild-type (WT) NifL and NifA under limiting nitrogen and oxygen (minus N). Typical WT activity in Miller Units was 1057 ± 84 (minus N), 32 ± 6.4 (plusN) and 325 ± 75 (plusO2), n=6.

A previously established two-plasmid system [20, 31, 32] was used to determine the *in vivo* activity of single amino acid substitutions in the *A. vinelandii* NifL and NifA proteins expressed in *Escherichia coli* strain ET8000 using a *nifH::lacZ* fusion as a reporter (Fig. 5D, E). Analysis of NifL variants (Fig. 5D) revealed that in all cases, amino acid substitutions increased NifA activity, as the reported β-galactosidase activity was always higher in the mutants compared to the wild-type under limiting nitrogen conditions (-N). A similar trend of increased relief of NifA inhibition, and therefore altered ability of NifL to interact with NifA, was observed under excess oxygen (+O2) conditions (Fig. 5D). However, in contrast, under excess nitrogen conditions (+N), several of the variants maintained the ability to inhibit NifA, at least partially, particularly in the case of alanine substitutions, indicating that the specific side chain is not crucial to the interaction under these conditions, in contrast to charge-chain substitutions at these positions that alleviated inhibition, presumably as a consequence of electrostatic repulsion (e.g. NifL substitutions at E298, R323 and E488). We attribute these differences in oxygen and nitrogen responses to the nature of the protein complexes formed in response to environmental signals, since alanine substitutions can evidently de-stabilize the NifL-NifA complex formed under oxidizing conditions, but not the GlnK-NifL-NifA complex formed under conditions of nitrogen excess. Quantitative western blot analysis of protein extracts derived from the same cultures used to measure β-galactosidase activity revealed that the substitutions did not significantly influence the expression levels of either NifL or NifA (Fig. S5).

Analysis of the influence of NifA substitutions on inhibition by NifgL was more challenging because the predicted contacts with NifL are located in the catalytic AAA+ domain (Fig 5, A, E). Nevertheless, we identified substitutions that impeded regulation by NifL and retained high NifA activity in the absence of NifL, indicating that catalysis was not impeded by these substitutions. In some cases, the activity of NifA variants followed a similar pattern to those observed for the NifL mutants described above. For the NifA residues K230, E356, E360, and E366, charge reversing substitutions led to complete escape from regulation by NifL in repressing conditions (+N and +O_2_), whereas substitutions to alanine resulted in repression only in the presence of excess nitrogen (+N) (Fig. 5E). Interestingly, some NifA variants, for example, P257A, R258D, and N373A were constitutively repressed by NifL even under permissive nitrogen-limiting conditions. This cannot be explained by a deficiency in NifA activity since these variants (indicated by asterisks in Fig. 5E) were fully active in the absence of NifL. Quantitative western blot analysis indicated that the activity patterns observed are unlikely to represent the influence of the amino acid substitutions on the stability of the proteins because, in almost all cases, we observed no differences in NifL and NifA stoichiometry (Fig. S5).

### Structure-guided mutagenesis of NifL and NifA enables unregulated expression of *nif* genes and ammonia secretion in *Azotobacter vinelandii*

Following up on our analysis of NifL and NifA mutant activities in a heterologous expression system (Fig. 5D, E), we further analyzed the nitrogen response of selected variants when expressed from their native genomic location in *A. vinelandii*, using a dual translational *PnifH::GFP::LUX* reporter fusion located at a neutral site (*algU*) in the genome [26] to quantify *nif* gene expression. Nine protein variants in *A. vinelandii*, were selected from those characterized in *E. coli,* together with the previously characterized NifA-E356K variant [26], and analyzed for nitrogen regulation of *nifH* expression and nitrogenase activity. Since escape from nitrogen feedback regulation is likely to result in ammonium excretion [26], a desirable trait for engineering more efficient diazotrophs, we also measured the ammonia concentration in culture supernatants (Fig. 6). In contrast to the native *nifLA* background strain (Av_646), most of the variants exhibited levels of *nifH* expression under excess nitrogen (+N, black bars) comparable to the levels of expression of the wild-type under limiting nitrogen conditions (-N, pink bars, Fig. 6A). Measurement of nitrogenase activities under the same conditions using the acetylene reduction assay revealed that the levels of *nifH* expression correlated well with the level of acetylene reduction activity observed under excess nitrogen conditions (+N, Fig. 6B). Most variants that ablated nitrogen feedback regulation and exhibited substantial nitrogenase activity in the presence of excess nitrogen, were also able to excrete ammonia in the millimolar range (Fig. 6C). Overall, these results in *A. vinelandii* correlate well with the nitrogen regulation data observed in the heterologous *E. coli* system. However, the NifA-K230E variant is an exception since it was blind to nitrogen source repression in the heterologous system (Fig. 5E) but failed to excrete ammonia in *A. vinelandii (*Fig. 6C). Quantitative western blot analysis revealed that the NifA-K230E variant is potentially unstable in *A. vinelandii*, thus providing an explanation for this discrepancy (Fig. 6D). All other variants did not exhibit significant changes in the relative abundances of NifL and NifA, reinforcing the conclusion that variant phenotypes reflect perturbations in surface interactions between both regulators.

**Figure 6.**
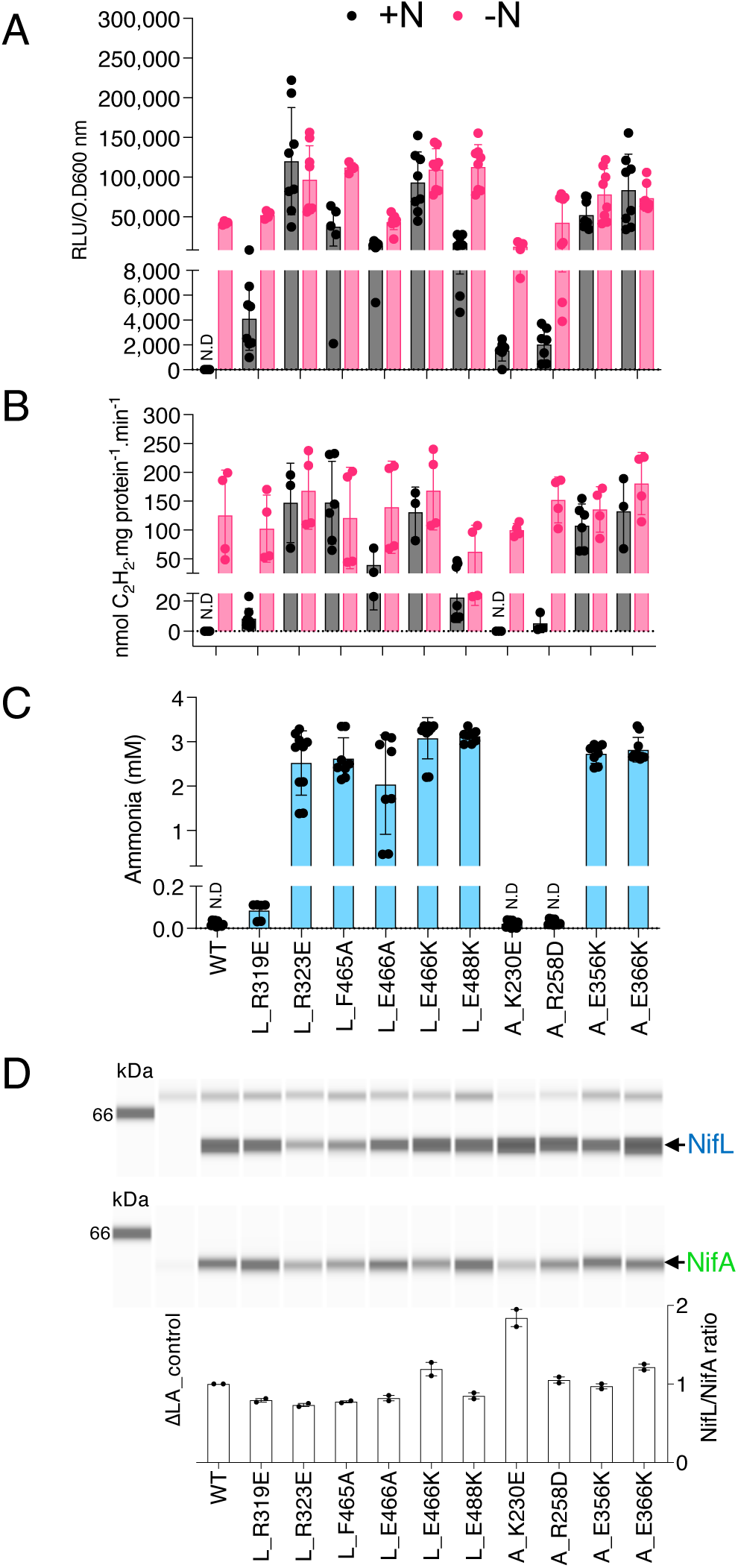
Characterization of NifL and NifA single amino acid substitutions in *Azotobacter vinelandii*. (A) Activity of a chromosomally integrated *PnifH::lux* fusion under excess (+N, grey) or limiting nitrogen (-N, pink) conditions in the wild type, strain Av_646, compared to various mutant strains. Activity is presented as relative luminescence units normalized by the cell density (RLU/O.D600nm). (B) *In vivo* nitrogenase activity. The activity was determined by the acetylene reduction assay using cultures grown under excess (+N, grey) or limiting nitrogen (-N, pink) conditions. (C) Ammonia quantification from the supernatant of cultures grown under limiting nitrogen conditions. (D) Quantitative immunodetection of NifL and NifA in the strains indicated.

## Discussion

In many cases, the activity of bacterial enhancing binding proteins (bEBPs) is tightly regulated by restricting the ability of the AAA+ domain protomers to assemble into a catalytically competent oligomer, typically a hexamer or heptamer (for extensive reviews, see [12, 33, 34]). Regulation of the AAA+ domain can be either exerted *in trans* through a partner regulatory protein or *in cis* through the bEBP N-terminal regulatory domain. For example, NtrC1 from *Aquifex aeolicus* [35] and NorR from *E. coli* [36] are negatively regulated *in cis* by their N-terminal regulatory REC and GAF domains, respectively. PspF from *E. coli* [37], a bEBP lacking an N-terminal regulatory domain, is negatively regulated *in trans* by interaction with its anti-activator partner, PspA. Interestingly, PspA does not interfere with the oligomerization of PspF. These proteins can form a complex comprised of six PspF monomers and six PspA monomers that can still interact with the RNA polymerase closed complex but are deficient in ATP hydrolysis [37]. Similarly, the regulation of AAA+ activity in NorR does not involve a change in the oligomerization state. The GAF domain, in hexameric DNA-bound NorR, negatively regulates the activity of the AAA+ domains by preventing access of the surface-exposed loops to σ54 of RNA polymerase. Upon nitric oxide binding, the GAF domain relieves repression of the AAA+ domain, enabling ATP hydrolysis [38].

In this study, we uncovered the molecular basis of the interaction between the inhibitory form of the anti-activator protein NifL and the bEBP NifA from *A. vinelandii*. The oxidized form of NifL sequesters NifA monomers into a tetrameric NifL2:NifA2: complex, facilitated by stochiometric levels of NifL and NifA expression. The binding of ADP to the GHKL domain, which induces conformational changes in the C-terminal region of NifL [20, 27, 30], is required to stabilize the complex and is likely to contribute to the extensive interaction surface involving both the GHKL and DH domains of NifL.

Although nucleotide binding by the AAA+ domain of NifA is unlikely to be impeded in the complex, ATP hydrolysis is prevented by the separation of NifA protomers to opposite sides of the NifL dimer. Formation of the catalytic site for ATP hydrolysis in bEBPs, requires interactions *in trans* between residues in adjacent AAA+ domain protomers, normally positioned in a “back-to-front” configuration in the hexamer (Fig. 7). A precedent for negative regulation of bEBPs *in trans* by protein partners is observed for the analogous tetramer formed between the FleQ bEBP and the anti-activator FleN from *Pseudomonas aeruginosa* (Fig. 7B). Similar to the NifL-NifA complex, the FleN dimer is sandwiched between the AAA+ domains of FleQ, and results in a conformational change in FleQ that prevents ATP binding and productive interactions with σ^54^ [39, 40] under certain conditions. In contrast to regulation via anti-activator-bEBP interactions, control of catalytic activity in the NtrC1 bEBP occurs through intramolecular interaction between the REC and AAA+ domains, which is regulated by phosphorylation. In the non-phosphorylated dimeric form of NtrC1, the AAA+ domains are held together by a REC domain dimer in a configuration that prevents catalysis. Phosphorylation is predicted to alter the conformation of the REC domain, leading to the formation of an active heptamer in which the AAA+ domains assemble in a “back-to-front” configuration that favors ATP hydrolysis (Fig. 7C).

**Figure 7.**
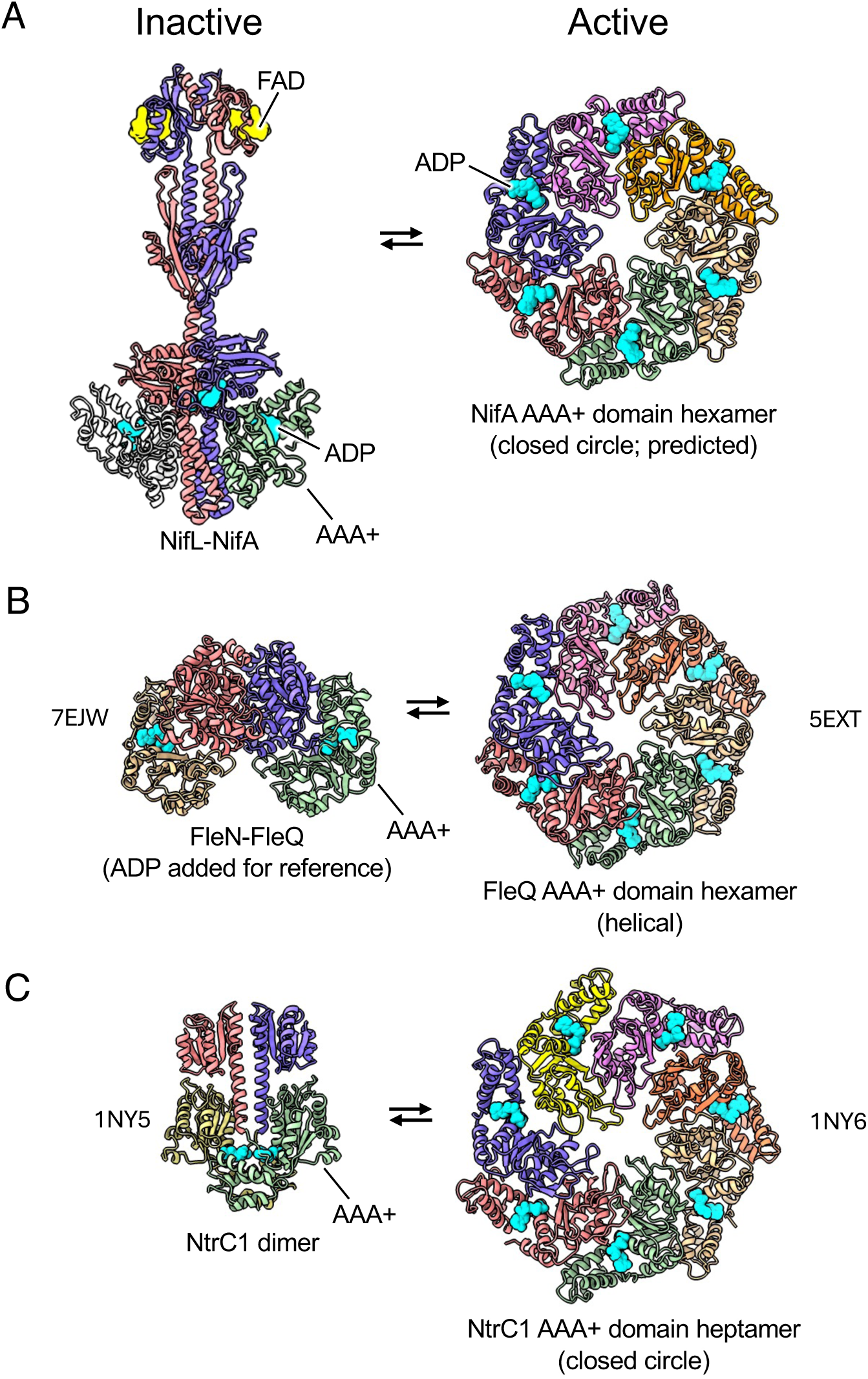
Comparison of inactive and active forms of σ^54^-dependent transcriptional activators. In the inactive forms, the AAA+ ATPase domains are kept apart and prevented from forming active homo-multimers. (A) In the cryo-EM structure of the NifL-NifA complex described herein, only one copy of the AAA+ domain of NifA is resolved, although AlphaFold3 predictions, place a second copy on the opposite side. We have added this second copy for comparison with the other structures (see also Fig. S2). There is no experimental structure of the active form of NifA, but a hexameric assembly of the AAA+ domain can be predicted by Alphafold3 with good confidence (ipTM = 0.68, pTM = 0.72). (B) In the FleN-FleQ complex (PDB code 7EJW) the FleQ AAA+ domains are separated by a central FleN homodimer (in the image shown, ADP molecules are added for reference purposes only – these were not present in the experimental structure). The activated FleQ AAA+ domains form a right-handed helical assembly, repeating every six protomers (PDB code 5EXT). (C) In the NtrC1 dimer (PDB code 1NY5), the C-terminal AAA+ domains (green and yellow) are kept apart by the N-terminal receiver domains (salmon and purple), and the activated AAA+ domains form a heptameric assembly (PDB code 1NY6).

In order for NifA to escape inhibition by NifL under conditions favorable for nitrogen fixation, conformational changes in both proteins are likely to be required. We have previously demonstrated that reduction of the FAD cofactor in the PAS1 domain results in a more kinked conformation in which the relative orientation of the GHKL and DH domains apparently changes to restrict access to NifA [20, 27]. Moreover, the GAF domain of NifA has an important role in enabling NifA to escape NifL inhibition via a conformational change induced by the binding of the ligand 2-oxoglutarate [32, 41]. In the cryo-EM map presented here, we could not observe density corresponding to the GAF domain in the full-length complex modeled by AlphaFold. This is potentially due to the high flexibility of the helical linker connecting the GAF domain to the AAA+ domain, which may have added to the challenge of fully resolving the structure of this complex to a higher resolution. Further attempts to stabilize this inhibitory complex by adding a third protein, GlnK, that contributes to nitrogen signaling, may stabilize the complex into a form more amenable to high-resolution cryo-EM in the future.

The structural similarity between NifL and histidine protein kinases (HPKs) raises questions concerning the evolution of these proteins and the role of their structurally conserved C-terminal GHKL and DH domains, which constitute the catalytic core of HPKs. The signal transduction mechanism of HPKs involves activation via rotation of the GHKL domain (otherwise known as the CA or catalytic domain). This enables either *cis* or *trans* autophosphorylation of the conserved histidine residue in the DHp domain and docking of the response regulator adjacent to the phosphorylated histidine to enable phosphotransfer to the response regulator. Substantial rotations of the GHKL domain and changes in the conformation of the DHp helices accompany the transition from the phosphotransferase state to the phosphatase state (Fig. S6). In contrast, the oxidised NifL-NifA complex is inherently more stable than the transient interactions between HPKs and response regulators and has a buried surface area of 1745 Å^2^, reflecting the extensive interactions between the GHKL and DH domains of NifL with NifA.

Our extensive functional analysis of single amino acid substitutions in the NifL-NifA interface not only uncovers the importance of specific amino acid chains for stabilization of the complexes formed between NifL and NifA but provides a blueprint for engineering control of nitrogen fixation in diverse diazotrophs, since the NifL-NifA interaction surface is likely conserved across a broad phylogenetic range of nitrogen-fixing proteobacteria. Structure-informed single amino acid changes confer advantages over harsher approaches, such as deleting *nifL*, since they enable controlled modulation of NifA activity, rather than unregulated runaway *nif* expression in *nifL*-null mutants that results in a severe growth phenotype or, in some cases, lethality [42, 43]. Further exploration of this approach based on refined control of NifL-NifA interactions may lead to the development of new approaches for engineered control of nitrogen fixation, thus making an important contribution to sustainable agriculture.

## Material and Methods

### Bacterial strains and growth conditions

All bacterial strains used in this study are listed in Table S1. *E. coli* strains were grown at 37°C in LB medium [44] for routine procedures or in NFDM [45] for β-galactosidase activity assays. Media for *E. coli* ST18 [46] was supplemented with 50 μg/mL of ALA (5-aminolevulinic acid) unless counter-selection was required. *A. vinelandii* was grown at 30°C and 250 rpm in NIL or MBB media as described previously [26, 32, 47]. Antibiotics were used as follows: carbenicillin 50 μg/mL (*E. coli*), chloramphenicol 15 μg/mL (*E. coli*), kanamycin 50 μg/mL (*E. coli*) and 2 μg/mL (*A. vinelandii*), trimethoprim 15 μg/mL (*E. coli*) and 90 μg/mL (*A. vinelandii*). Growth curves were performed on a 24-well microplate (Greiner-Bio one #662160) using a Biotek EON plate reader, as reported before [26]. The absorbances recorded at 600 nm were corrected to a 1 cm path length.

### Molecular Cloning

All the plasmids used in this study are listed in Table S2. General molecular biology techniques were performed according to established protocols [44]. Enzymatic isothermal assembly [48] was performed with the NEBuilder HiFi DNA Assembly Master Mix (NEB #E2621), whilst site-direct mutagenesis was performed using Q5® Site-Directed Mutagenesis Kit (NEB #E0554) according to the manufacturer’s protocols. High-fidelity DNA polymerase and restriction enzymes were provided by New England Biolabs. DNA purification was performed using commercially available kits provided by Macherey-Nagel. Sanger DNA sequencing was conducted by Genewiz-Azenta Life Sciences. IDT Technologies conducted oligonucleotide synthesis, whilst gene synthesis was performed by Genscript or Twist Bioscience.

### Construction of mutants in *A. vinelandii*

Detailed procedures for *A. vinelandii* genetic manipulation have been published elsewhere [26, 49, 50]. An *A. vinelandii* strain (Av_2302) carrying a *PnifH::GFP-LUX* translational fusion in addition to a *nifLA* deletion (*nif-*) was used to re-insert *nifL* and *nifA* mutant alleles and enable selection of active mutant variants via recovery of the *nif+* phenotype (ability to grow diazotrophically in media lacking fixed nitrogen). To construct Av_2302, we first transformed competent *A. vinelandii* DJ cells prepared as described earlier [49] with the plasmid pKS1909 linearized with ScaI. A recombinant colony carrying the *PnifH::GFP-LUX* translational fusion was selected in media containing trimethoprim and named Av_KS425. Subsequently, Av_KS425 was conjugated with *E. coli* ST18 carrying pMB2245 to generate a *nifLA* deletion. Single crossover recombinants were selected in a medium containing glucose as a carbon source and kanamycin. After confirmation of plasmid integration by PCR, double crossovers were selected in media containing sucrose and ammonium acetate as nitrogen source. Deletion of *nifLA* was confirmed by PCR and by the lack of growth in media without added fixed nitrogen (*nif-*). Subsequent transfer of *nifL* and *nifA* point mutants to *A. vinelandii* was achieved by conjugating pK18mobsacB derived vectors (see Table S2) encoding the desired mutant *nifLA* wild-type or mutant operons into the strain Av_2302. After selection of single crossovers as above, double crossovers were selected in media containing sucrose without any added nitrogen. Mutants were selected solely on their ability to restore diazotrophy (growth using atmospheric N_2_ as the sole nitrogen source). PCR fragments obtained with oligonucleotides flanking the *nifLA* region were sequenced to confirm the integration of the desired mutation. All strains are listed in Table S1.

### β-galactosidase activity

β-galactosidase activity assays were performed as described previously [51] using growth conditions previously established [20, 31, 32], except that 6 mL screw-capped bijou universals were filled to the brim to enable limiting oxygen conditions [26, 52]. To facilitate overall activity comparison across all mutants tested, the activity is presented as *log2FC* respective to the activity of the wild-type NifL-NifA pair expressed from pPR34 under limiting oxygen and nitrogen (-N, maximum activation conditions). Typical mean activities in Miller Units were 1057 ± 84 (-N), 32 ± 6.4 (+N), and 325 ± 75 (+O_2_), n=6. Note that for the wild-type, the ratio between each replicate with the mean under -N is approximately 1, and therefore, its *log2FC* is approximately 0 (orange in the scale bar). *log2FC*>0 (green) indicates hyperactivation, whilst *log2FC*<0 (red) indicate repression.

### Luminescence reporter assays

To measure the expression of nitrogenase, we used a chromosomally integrated translational *PnifH::GFP::LUX* fusion constructed as described above. *A. vinelandii* strains were grown exactly as described for the nitrogenase activity assays. After incubation, 200 μL of cells were transferred to a 96-well plate black plate with a clear bottom to avoid signal bleeding across wells and reflection interferences. The luminescence was quantified using a BMG ClarioStar plate reader set to a focal height of 9.7 mm and a monochromator gain set to 3600. Before data acquisition, the full plate was scanned, and data was acquired with a 70% gain adjustment relative to the highest signal detected in each plate. The luminescence output reported as relative luminescence units (RLU) was normalized by O.D_600nm_ of the same culture aliquots measured in a replicated transparent 96-well plate. Background signals from media only and strains encoding the LUX fusion in a double *nifLA* mutant background (strain Av_2302) or not encoding the *PnifH::GFP::LUX* fusion (Av_DJ) were subtracted from the luminescence derived from the test samples. The luminescence signal for background controls and assay validation samples (strains Av_646, wild type and Av_614, NifA_E356K) were also visualized using a Berthold NightOwl camera (Fig. S7) for additional validation.

### Nitrogenase activity

*In vivo* nitrogenase activity in *A. vinelandii* was measured by the acetylene (C_2_H_2_) reduction assay [53, 54] in cultures growing in NIL liquid media as indicated below. After incubation of cells with acetylene for 2 to 3 hours, ethylene (C_2_H_4_) was quantified using an Agilent GC 8860 gas chromatograph equipped with a HayeSepN 80/100 (Agilent #G3591-80037) column. The injector was kept at 150°C, the oven at 140°C, whilst the FID detector was set at 250°C. Air flow from a zero-air generator with an integrated compressor (Chromalytic model #ZA2A) was set at 400 mL.min^-1^, hydrogen fuel was set at 30 mL.min^-1^, hydrogen carrier flow was set at 20 mL.min^-1^, and nitrogen at 25 mL.min^-1^ was used as make-up flow. The instrument was calibrated using ethylene derived from the chemical decomposition of ethephon (Sigma #C0143) in a 10 mM Na_2_HPO_4_ pH = 10.7, as described previously [55]. An ethylene calibration ranging from 10 – 200 nmol was interpolated using the Agilent OpenLab software and automatically applied to a manual acquisition method with the parameters described above. As *A. vinelandii* is able to fix nitrogen in air (21 % O_2_), 10 mL cultures were grown in NIL media as above in 50 mL conical flasks in an open atmosphere at 250 rpm until the desired O.D_600nm_ was reached. Immediately before the acetylene reduction assay, the flask was stopped with rubber septa (Suba-Seal® n°37), 10% acetylene was injected, and the ethylene formed was analyzed as described. Nitrogenase activity is reported as nmol of C_2_H_4_.mg protein^-1^.min^-1^.

### Quantification of ammonia

Ammonia was quantified as described in [26] using an indophenol-blue-based method modified from [56, 57].

### Protein expression and purification

Proteins were purified from 2-4 liters of *E. coli* Nico21 (DE3) grown at 30°C, 250 rpm. Once the cultures reached an OD_600 nm_ of around 0.5–0.6, protein overexpression was induced for 3 hours by adding 0.5 mM IPTG. The cells were harvested by centrifugation, and the cell paste was resuspended in Buffer A_STH (20 mM HEPES pH 7.5, 150 mM NaCl, 25 mM MgCl_2_, 2.5% Glycerol, 1 mM ADP) or Buffer A_HT (20 mM HEPES pH 7.5, 300 mM NaCl, 2.5% Glycerol) and frozen at -20°C. For Av-Strep-tag-NifL (or Av-Strep-tag-NifL_269-519_) and its co-purifications with Av-NifA-6xHis (or Av-NifA_188-455_-6xHis), the Buffer A_STH was used, whilst Buffer A_HT was used to purify Av-SUMO-NifA-6xHis. The next day, after thawing, the cells were disrupted by 3-4 passages through an Avestin EmulsiFlex-B15 cell homogenizer at an air pressure of 40 psi (12,000 psi homogenizing pressure) and immediately centrifuged at 37,000 g at 4°C for 30 minutes. The supernatant was filtered through a 0.22 μM membrane and then loaded onto a 5 mL HiTrap StrepTrap XT column (Cytiva #29401320) previously equilibrated with Buffer A_STH for Av-Strep-NifL. The supernatant from Av-SUMO-NifA-6xHis expression was loaded onto a 1 mL HisTrap Excel (Cytiva #29048586) equilibrated with Buffer A_HT. Av-Strep-NifL on its own or in complex with NifA variants was eluted in a step gradient with 100% Buffer B_STH (Buffer A_STH with added 15 mM biotin). Av-SUMO-NifA-6xHis was eluted in a gradient elution over 10 column volumes with Buffer B_HT (Buffer A_HT amended with 0.5 M imidazole). For co-purification of Av-Strep-tag-NifL (or Av-Strep-tag-NifL_269-519_) with Av-NifA-6xHis (or Av-NifA_188-455_-6xHis) fractions from the first chromatographic step were pooled, the salt concentration was raised to 300 mM NaCl (from a 5M stock solution) and finally concentrated using a 50 kDa cut-off Amicon Ultra-15 centrifugal filter (Merck #UFC905024). To ensure maximum homogeneity, the concentrated fractions were further purified by size exclusion chromatography using a Superdex 200 increase 10/300 GL (Cytiva #28990944) and eluted in SEC_300 Buffer (20 mM HEPES pH 7.5, 300 mM NaCl, 2.5 mM MgCl_2_, 1 mM ADP, 1 mM TCEP). All chromatography steps were performed using an AKTA pure FPLC system (Cytiva).

### Mass Photometry

The oligomeric state of purified protein complexes was assessed by mass photometry [58] using the OneMP mass photometer (Refeyn) instrument following the manufacturer’s instructions. Samples were prepared in PBS (8 mM Na_2_HPO_4_, 2 mM KH_2_PO_4_, 137 mM NaCl, 2.7 mM KCl, pH 7.4) amended with 5 mM MgCl_2_ and 1 mM ADP. Purified complexes were normalised to approximately 1000 nM, and then 1 uL was added to a 9 μL buffer droplet in the instrument objective slide. The instrument was calibrated by interpolating the ratiometric contrast signal against the molecular weight (kDa) of a set of calibrants as follows: β-amylase (56, 112, and 224 kDa), thyroglobulin (670 kDa), and urease (90.7, 272 and 545 kDa) also diluted to the same molar rage in the PBS Buffer.

### Automated western blots

Immunodetection of NifL and NifA from purified protein fractions or *E. coli* ET8000 and *A. vinelandii* whole cell extracts was performed using polyclonal antisera (Biosynth Laboratories) raised against Av-Strep-NifL or Av-SUMO-NifA-6xHis purified as described. Quantitative western blots were performed using the WES automated system (Protein Simple #004-600). Approximately 0.2 µg of purified protein diluted in 0.1X WES sample buffer was mixed with 1X Fluorescent Master Mix from the EZ Standard Pack (#PS-ST011EZ) in a final volume of 6 μL and loaded into a 12-230 kDa capillary cartridge (#SM W002) as recommended by the manufacturer. Whole-cell extracts were prepared by collecting the cell pellet from 0.8 mL cultures (*E. coli* O.D_600nm_: 0.7 – 1.2 and *A. vinelandi* O.D_600nm_: 0.2) and resuspending in 1-2X Novex Tris-Glycine SDS Sample Buffer (Thermo Scientific #LC2676). 1-2 μL of the above extract was diluted in 0.1X WES sample buffer and mixed with 1X Fluorescent Master Mix as described earlier. To ensure proper molecular weight estimation from whole-cell extract samples, the WES biotinylated MW ladder was diluted in a buffer containing the same final concentration of the Novex Tris-Glycine SDS Sample Buffer present in the final dilution of the whole-cell extract sample.

### Protein structure modeling

Structural modeling was performed with AlphaFold2-Multimer [59], as implemented in ColabFold [60], and the AlphaFold3 server [61]. A full list of protein sequences used for modeling is available in Table S3. The starting model for refinement of the cryo-EM structure was generated using AlphaFold3, which allowed the incorporation of biologically-relevant ligands. This was successful in inserting FAD into each NifL PAS1 domain, and ADP in each NifL GHKL domain and in each NifA AAA+ domain. In all cases, the placement of these ligands was consistent with their placement in related experimental structures. Models were visualized and images were generated in UCSF ChimeraX v1.8 [62].

### Cryo-EM sample preparation and data collection

Freshly purified protein complex samples were diluted to approximately 4-5 μM in SEC buffer. Using a Vitrobot Mark IV (Thermo Fisher Scientific) at 4 °C and 100% relative humidity, 3.5 μL of purified protein sample was applied to a r2/2 holey carbon 300 mesh Cu grid (Quantifoil Micro Tools), which had been glow discharged for 60 seconds at 8 mA using an ACE 200 (Leica Microsystems). Twenty seconds after sample application to the grid, the grid was blotted for 4 seconds with a blotting force of 8 and then plunged into liquid ethane. Cryo-EM data collection was performed at the Electron BioImaging Centre (eBIC; Harwell, UK) on a Titan Krios transmission electron microscope (“Krios 2”, Thermo Fisher Scientific) operated at 300 keV with a K3 Summit direct electron detector (Gatan). Movies, comprised of 50 frames each, were collected using EPU software v3.5.1, with a nominal magnification of 105kX, pixel size of 0.831 Å, defocus range of −0.7 to −2.2 μm total dose of 50 e^−^/Å^2^. Further data collection details are summarized in Table S4.

### Cryo-EM data analysis

The particles had a tendency to aggregate, were prone to orientational bias and because they were long and thin, they were difficult to distinguish from the background. Hence, data analysis presented a significant challenge. Single particle reconstruction was performed using CryoSPARC v4.5.3. Manual particle picking was not possible; thus particles were picked from the whole dataset using an elliptical blob picker. From these, a set of eight templates were produced after 2D classification, which were then used for template-based picking and, after one round of further 2D classification, yielded a stack of ∼2.2m particles. The classes revealed an elongated particle with high-resolution features, although there was also some blurring at the extremities. The best 2D-classes were fed into an ab initio reconstruction job with four classes requested, and all outputs were passed directly into a heterogeneous refinement job. This gave two classes that showed features consistent with AlphaFold models of the NifL-NifA complex. Both of these classes were independently taken forward for further classification and refinement using the same workflow, but one resulted in a significantly less complete reconstruction and was thus abandoned. The particles comprising the superior class from the heterogeneous refinement job yielded a stack of 743k particles after re-extraction to 1.48 Å/pixel. Following 3D classification into four classes, 209k particles from the resultant best class were passed to a final non-uniform refinement job, which gave an average resolution of 6.45 Å and, despite a clear preferential orientation, there were no significant gaps in the viewing angle coverage. The processing is summarized in Fig. S1. The reconstruction was clearly asymmetric, in contrast to the prediction of a two-fold symmetric particle by AlphaFold3. Nevertheless, there was good agreement between the NifL subunits and the cryo-EM density, which could be readily superposed in ChimeraX, with the density accounting for the majority of the sequence for the two NifL subunits (residues 21-517). There was a complete absence of density for one of the NifA subunits, even when the map contour level was lowered considerably, and there was significantly less density for the other NifA subunit compared to that which would be expected from the AlphaFold3 model. Closer inspection indicated that only the AAA+ domain was present (residues 210-458).

The top-ranked AlphaFold3 model (incorporating the expected ligands FAD and ADP) was used as a starting point for model building and refinement. A heterotrimer, consisting of two copies of NifL and one copy of the NifA AAA+ domain, was initially docked as a rigid body into the cryo-EM density in ChimeraX, and then real space refinement was performed in Coot using “self-restraints” to preserve the local stereochemistry of the starting model. This was then subjected to real space refinement in PHENIX, with the imposition of Ramachandran restraints as well as restraints to the input model. The latter restraints were subsequently turned off for a final PHENIX real space refinement job after adding hydrogens using phenix reduce. Compared to the rigid body docked AlphaFold3 model, the final model had rmsd values of 2.91, 3.05 and 2.16 Å for the two NifL subunits and the single NifA subunit, respectively, and an overall rmsd of 3.48 Å. Parameters of the final model are summarized in Table S5.

## Supporting information

BuenoBatista_etal_2015_SI

## Acknowledgments

We are grateful to all members of the laboratory support team at the John Innes Centre for their excellent technical assistance. M.B.B was supported by the UKRI-BBSRC (grant BB/W009986/1). M. W. W was funded by the BBSRC Institute Strategic Programme BRiC (grant BB/X01102X/1) and UKRI Future Leaders Fellowship (MR/X033481/1). D.G. is a Wellcome Trust Sir Henry Dale Fellow (221868/Z/20/Z); work in his lab is supported by the BBSRC Institute Strategic Programme HBio (grant BB/X01097X/1). J.W.P was supported by the US DOE, Office of Science, Office of Basic Energy Sciences (DE-SC0018143). R.D. was supported by the UKRI-BBSRC (grant BBS/E/J/000PR9797) and by the Royal Society grant ICA\R1\180088. The John Innes Centre Bioimaging Platform is supported by the UKRI-BBSRC (grant BB/CCG2240/1). M.B.B. received cryo-EM training from Astbury Structural Biology Centre, University of Leeds, through their Wellcome/MRC funded program (218785/Z/19/Z). Yehuda Halfon (Astbury Structural Biology Centre) is acknowledged for cryo-EM sample handling and training. Katherine Stewart is acknowledged for gifting plasmids pKS1901 and pKS1909. Julia Mundy and Abbas Maqbool are acknowledged for their technical assistance. We acknowledge Diamond for access and support of the cryo-EM facilities at the UK National Electron Bio-Imaging Centre (eBIC) under proposal BI34108.

## Author contributions

M.B.B. designed, performed research, acquired funding, managed the project, and wrote the manuscript with input from all authors. J.R. performed cryo-EM sample screenings and managed data acquisition. M.W.W., D.G., and D.L. designed cryo-EM sample preparation, data acquisition, and analysis. J.W.P. and R.D. structural model interpretation and manuscript writing.

## Data availability statement

All data needed to evaluate the conclusions in the paper are present in the main text or the Supporting Information. The NifL-NifA 2:1 coordinates have been submitted to the Protein Data Bank (https://www.rcsb.org/) with PDB ID 9QQ6. Corresponding EM maps have been submitted to the Electron Microscopy Data Bank (https://www.ebi.ac.uk/pdbe/emdb/) with ID EMD-53294. Research materials supporting this study are available from the corresponding authors.

