## Supplementary material for "Structural analysis of the NifL-NifA complex reveals the molecular basis of anti-activation of nitrogen fixation gene expression in *Azotobacter vinelandii*": BuenoBatista_etal_2015_SI

### Supplementary Figures

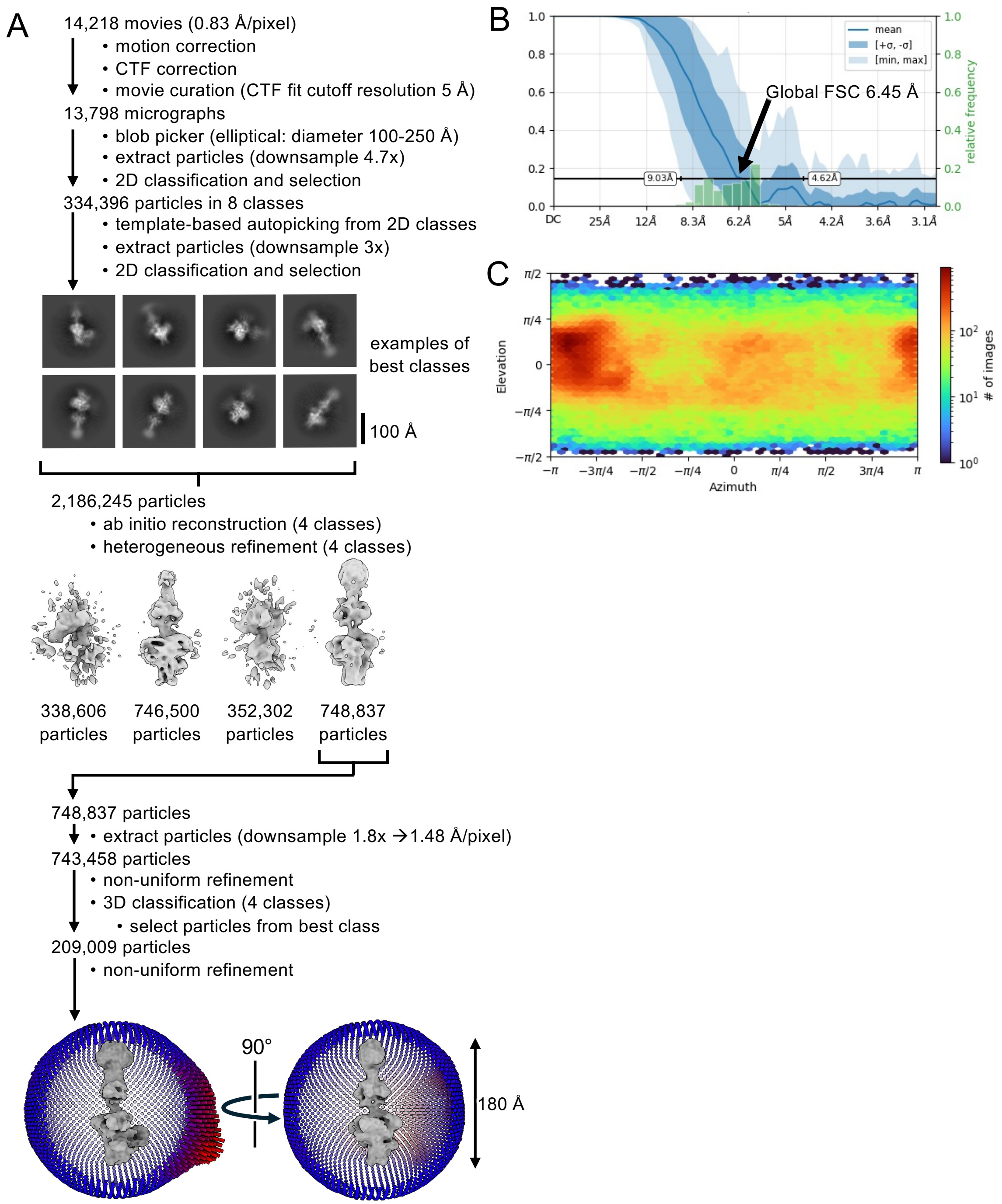


**Fig. S1.** (A) CryoSPARC single-particle analysis workflow for the NifL-NifA complex. The distribution of viewing angles (bottom) displayed on orthogonal views of the final reconstruction show a full coverage, but with a very distinct orientation bias highlighted by the red projections in the left-hand image. In both, the front views have been hidden for clarity by clipping the images. (B) Fourier Shell Correlations are plotted against spatial frequency, where the global values are represented by the dark blue line, with the variation and distribution in conical FSC values indicated by the blue shading and green histograms respectively. (C) Angular distribution plot of particle views.

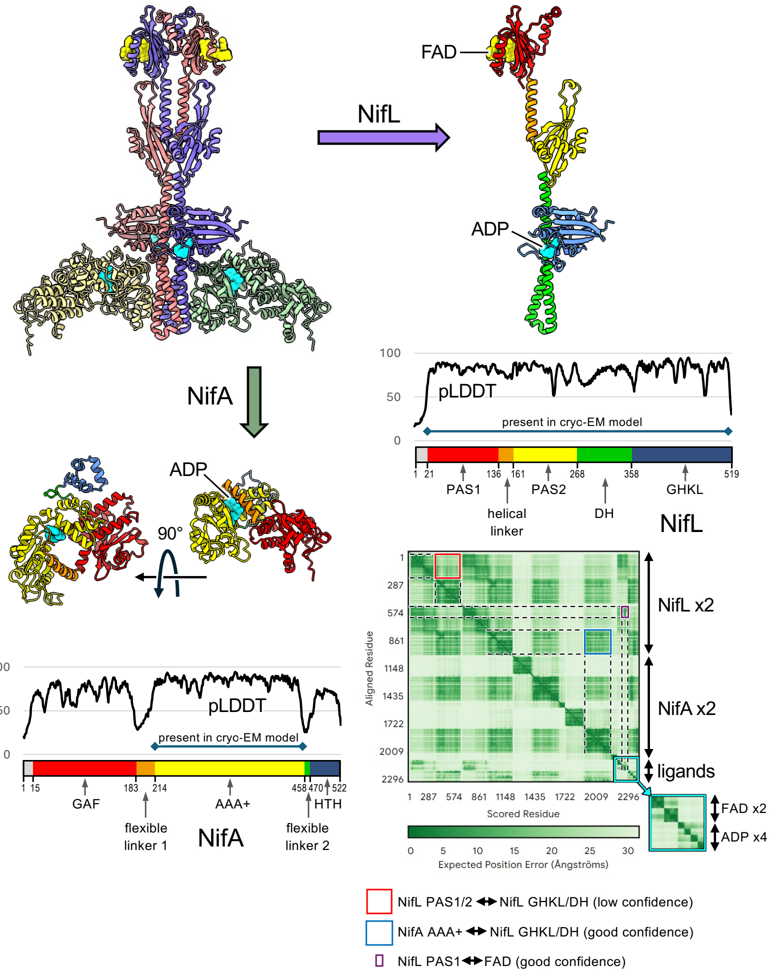


**Fig. S2.** AlphaFold3 prediction of the full-length NifL-NifA heterotetrameric complex structure. The model (top left) comprises two copies of NifL (salmon and purple cartoons) and two copies of NifA (green and pale-yellow cartoons) and displays two-fold symmetry about a vertical central axis. The domain topology is highlighted in the single separated subunits which are coloured according to their respective schemes in the panels below each. Also plotted in these panels are the per residue pLDDT scores indicating the local quality of the predictions. These clearly delineate the domain boundaries for NifA in particular as the scores are significantly lower for the short linker regions. The PAE plot (bottom right) shows the confidence in the relative placement of pairs of residues in the model. The individual domains are highlighted as distinct folded units by the darker green blocks along the diagonal of the plot. Off-diagonal blocks indicate the confidence in the relative placement of pairs of domains. Note that there is only off-diagonal pale green colouration for the regions corresponding to the placement of the NifL PAS domains with respect to the GHKL and DH domains (red box), indicating low confidence in their relative placement; this could be due to flexibility at the constriction between the upper and lower halves of the complex. By contrast, the AAA+ domain of NifA shows off-diagonal dark green blocks (blue box) indicating confidence in its placement relative to the NifL GHKL and DH domains. Additionally, smaller off-diagonal darker green blocks indicate good confidence in the placement of the ligands in their respective domains, for example, the placement of FAD in the NifL PAS1 domain (purple box). Overall, the predicted structure provides a testable model for investigating the structure-function relationships of the NifL-NifA complex. However, the pTM score of 0.55 indicates that the overall structure is likely to be similar to the true structures, whilst the ipTM score of 0.52 indicates relatively low confidence in the placement of the subunits within the complex.

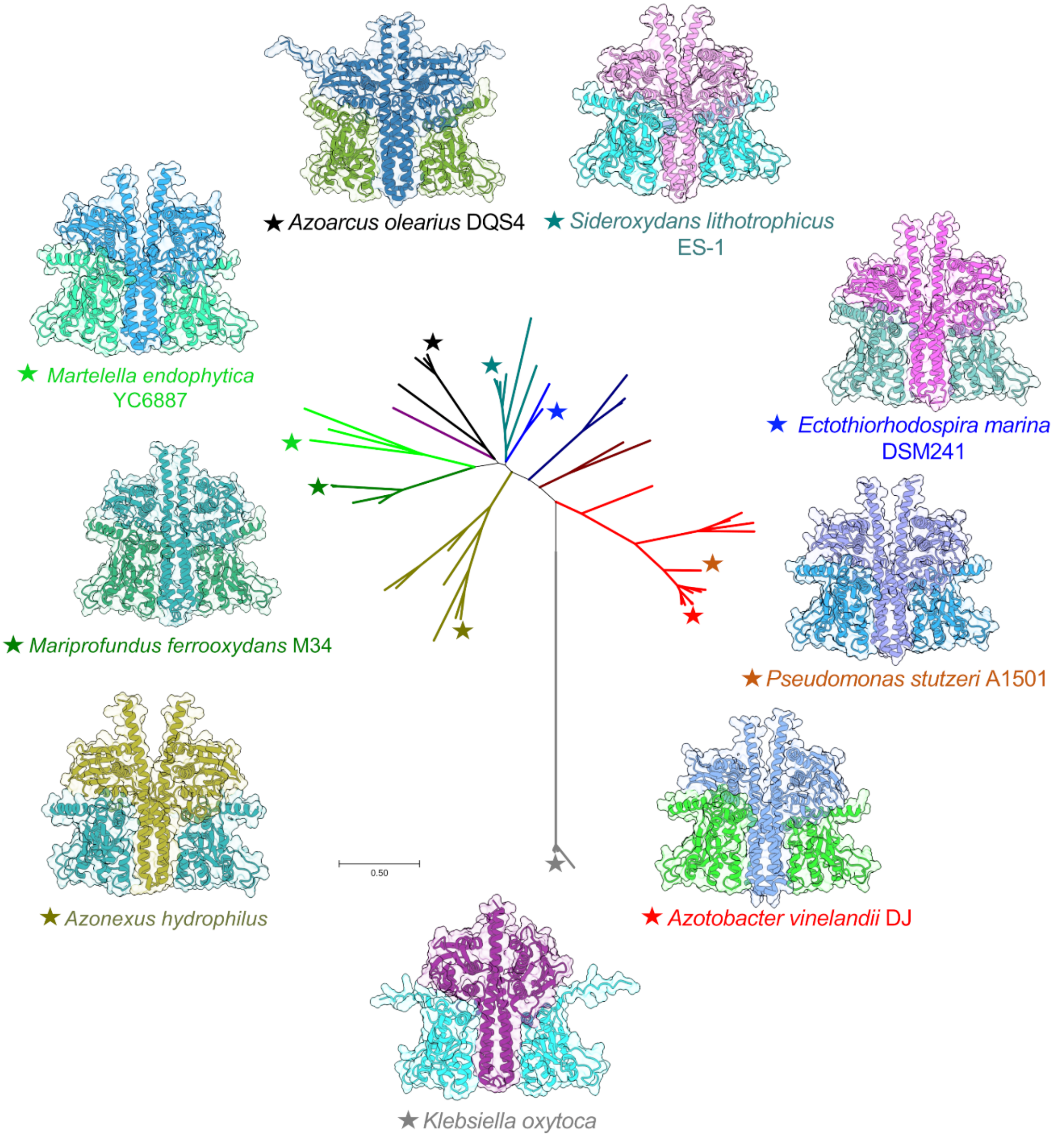


**Fig. S3.** Structural modeling of truncated NifL-NifA pairs from different bacterial species distributed across the phylogenetic spectrum in Proteobacteria. The phylogenetic tree in the center was inferred from the analysis of 60 NifL amino acid sequences as described previously (1). One or more NifL-NifA pairs from representative species from each major color-coded clade, was chosen for Alphafold2 multimer modeling as described in the material and methods. The amino acid sequences used for modeling are available in Table S4.

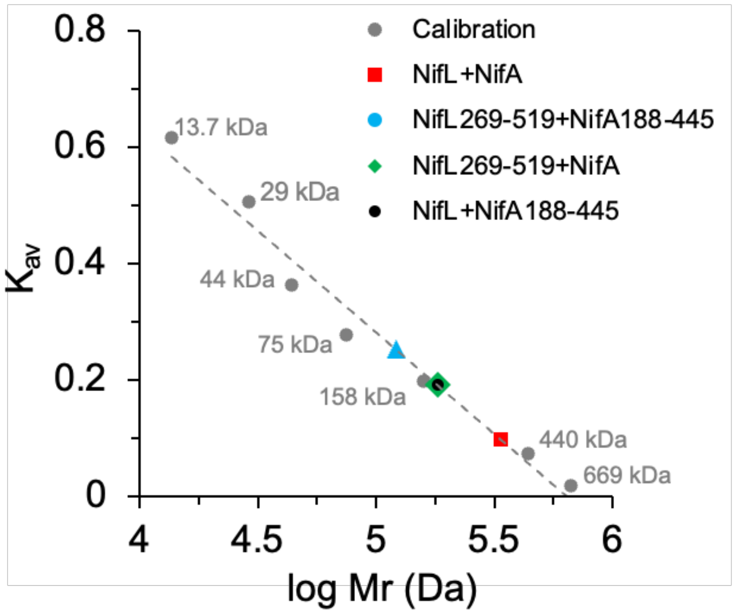


**Fig. S4.** Size exclusion chromatography profile of various purified NifL-NifA complexes compared to the molecular weight calibration standards (grey circles) resolved in a Superdex 200 increase 10/300 GL (Cytiva #28990944) column as described in the materials and methods. Red square: NifL-NifA, blue circle: $NifL_{269\text{-}519}$-$NifA_{188\text{-}445}$, green diamond: $NifL_{269\text{-}519}$-NifA and black circle: NifL-$NifA_{188\text{-}445}$. The standards used were as follows: Ribonuclease A (13.7 kDa), Carbonic anhydrase (29 kDa), Ovalbumin (44 kDa), Conalbumin (75 kDa), Aldolase (158 kDa), Ferritin (440 kDa) and Thyroglobulin (669 kDa).

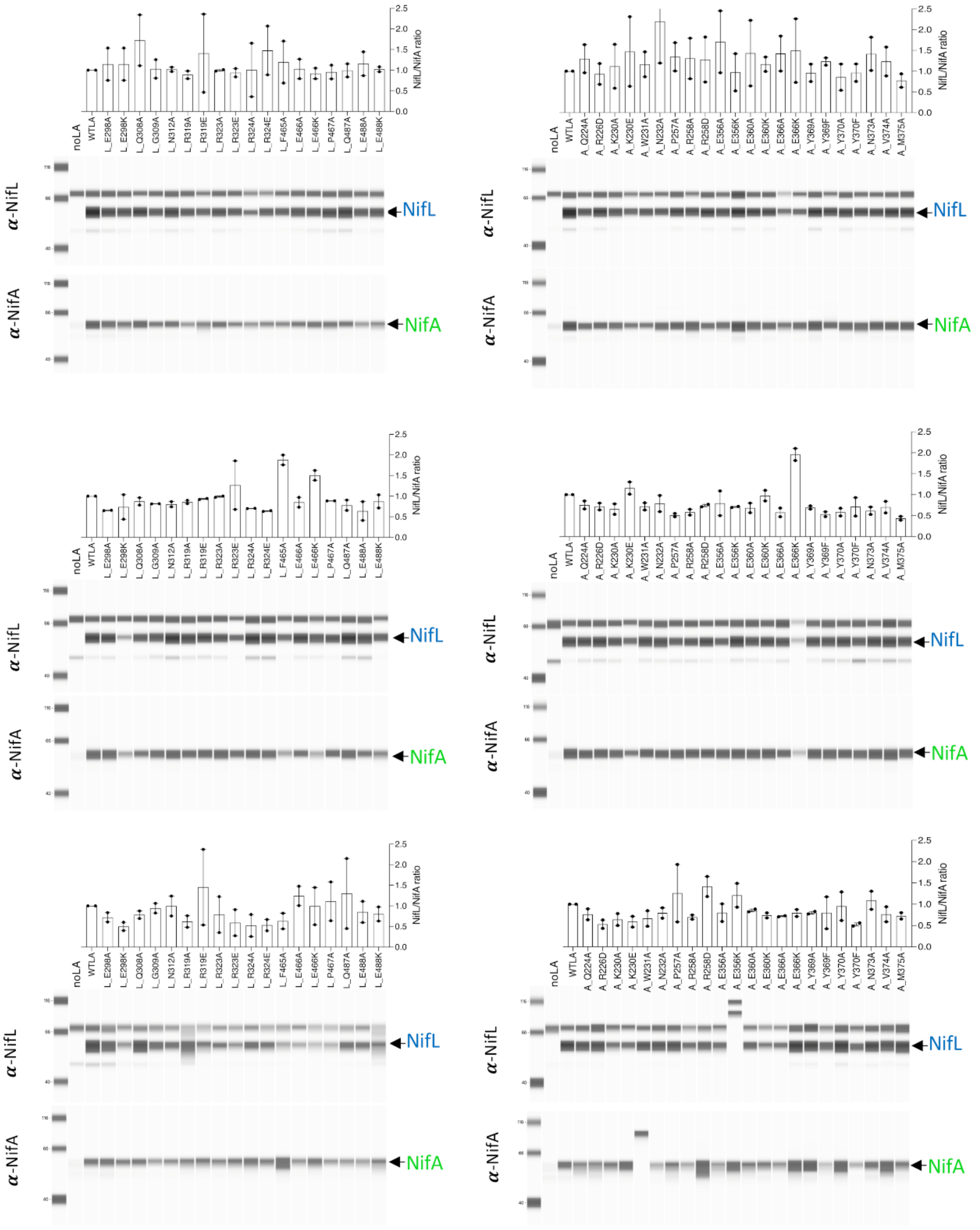


**Fig. S5.** Automated western blots used to quantify NifL/NifA ratios. Protein extracts were prepared from the same cultures used for activity determination in Figures 3D and 3E. Analysis of NifL point mutants is presented on the left side, whilst NifA mutants are presented on the right side. Conditions are as follows: top, limiting nitrogen (-N); middle, excess nitrogen (+N); bottom, excess oxygen $(+O_2)$.

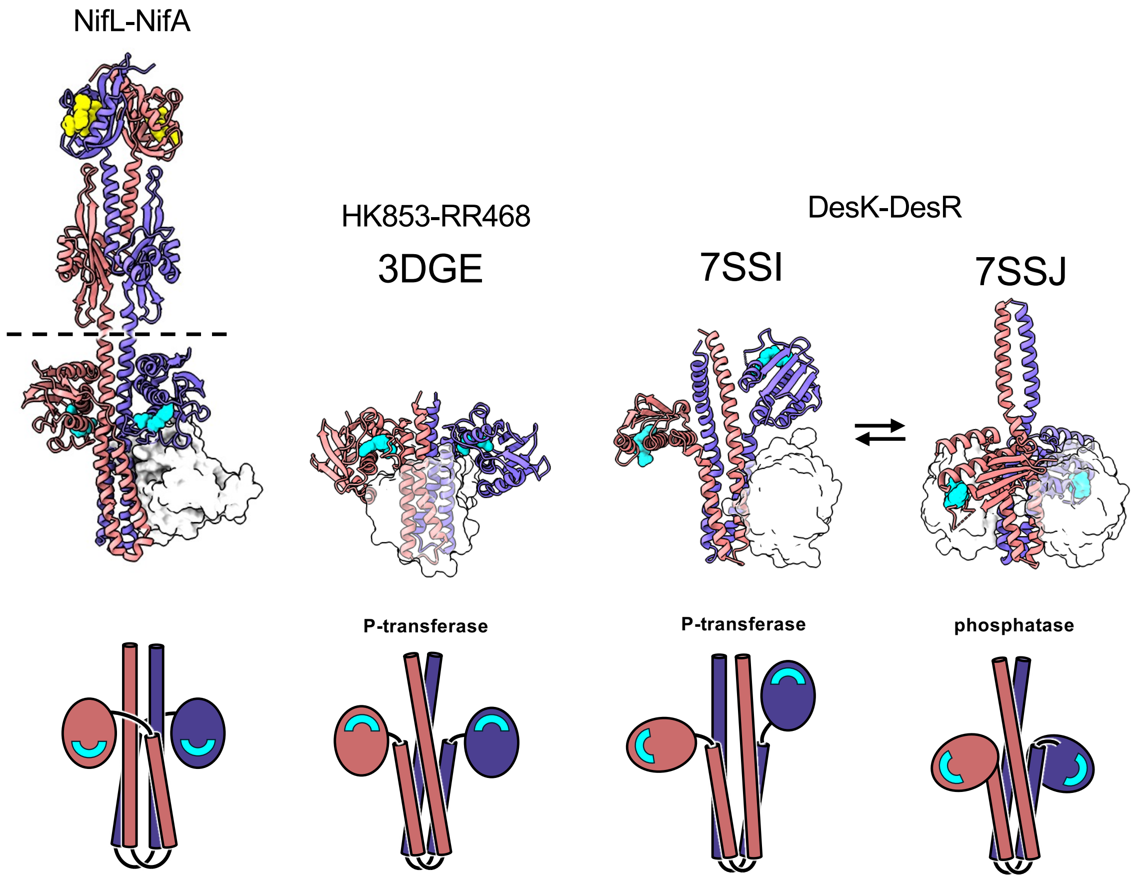


**Fig. S6.** Comparison of GHKL and DH domain topologies. The NifL GHKL and DH domains are topologically similar to structurally characterised members of the histidine protein kinase family in which the four helix bundle formed by the DHp domain in the HPK dimer interacts with the catalytic GHKL domain responsible for autophosphorylation of the conserved histidine residue.A common feature in HPKs is the switch in conformation between assymetic and symmetric states as illustrated by the conformational transition of DesK from the phosphotransferase to the phosphatase state as shown here. The interacting partners for NifL (AAA+ domain of NifA) and for HPKs (receiver (REC) domains of response regulators) are shown in spacefill. ATP/ADP binding sites are indicated in cyan.

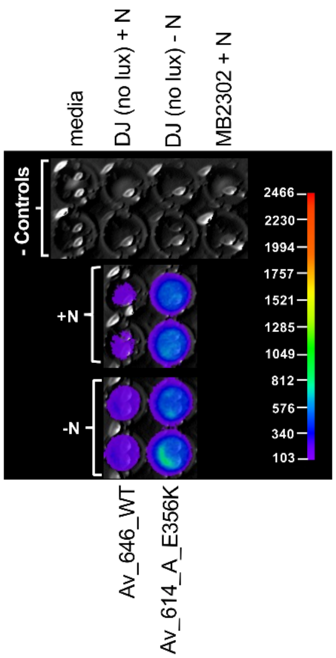


**Fig. S7.** Raw luminescence data validating our *PnifH::GFP::LUX* fusion used to quantify expression of *nif* genes in liquid cultures. The image was obtained using a Berthold NightOwl camera as described in materials and methods. In the top two rows (-controls) no background luminescence can be observed in a strain not encoding the fusion, DJ, or in a strain encoding the fusion but lacking both *nifL* and *nifA*. In the middle row, some background luminescence can be observed under excess nitrogen (+N) for the strain Av_646 (wild type), which is much lower than the luminescence observed for the strain Av_614 (NifA_E356K). In the bottom row, the luminescence under limiting nitrogen (-N) for both strains is shown. For the activities presented in Figure 3, any background luminescence in +N for the Av_646 (wild type) strain was subtracted from the remaining test strains.

**Table S1.** Strains used in this study.

| Strain | Description | Reference |
|---|---|---|
| *E. coli* | | |
| ET8000 | ET8000 robust E. coli strain for B-gal assays | (2) |
| ST18 | *E. coli* S17λpir Δ*hemA.* Used for conjugations | (3) |
| NEB 5-alpha | Derived from DH5α. T1 phage resistant and *endA* deficient. Used for cloning and plasmid maintenance | NEB |
| Nico21 (DE3) | *can::CBD fhuA2 [lon] ompT gal (λ DE3) [dcm] arnA::CBD slyD::CBD glmS6Ala ΔhsdS λ DE3 = λ sBamHIo ΔEcoRI-B int::(lacI::PlacUV5::T7 gene1) i21 Δnin5* | NEB |
| *A. vinelandii* | | |
| DJ | ATCC BAA-1303. High frequency transforming variant of *A. vinelandii* UW generated in 1984 by Dennis Dean | (4) |
| Av_4-25 | Derived from DJ. *PnifH::GFP-lux* inserted into the *algU* locus. $Tp^R$ | This study |
| MB2302 | Derived from Av_4-25 carrying a deletion of the *nifLA* operon. $Nif^-$ | This study |
| Av_646 | Derived from MB2302, selected for reconstitution of Nif+ with WT *nifLA* | This study |
| Av_673 | Derived from MB2302, selected for reconstitution of Nif+ with *nifL_R319E* and WT *nifA* | This study |
| Av_622 | Derived from MB2302, selected for reconstitution of Nif+ with *nifL_R323E* and WT *nifA* | This study |
| Av_663 | Derived from MB2302, selected for reconstitution of Nif+ with *nifL_F465A* and WT *nifA* | This study |
| Av_716 | Derived from MB2302, selected for reconstitution of Nif+ with *nifL_E466A* and WT *nifA* | This study |
| Av_658 | Derived from MB2302, selected for reconstitution of Nif+ with *nifL_E466K* and WT *nifA* | This study |
| Av_632 | Derived from MB2302, selected for reconstitution of Nif+ with *nifL_E488K* and WT *nifA* | This study |
| Av_686 | Derived from MB2302, selected for reconstitution of Nif+ with WT *nifL* and *nifA_K230E* | This study |
| Av_718 | Derived from MB2302, selected for reconstitution of Nif+ with WT *nifL* and *nifA_R258D* | This study |
| Av_614 | Derived from MB2302, selected for reconstitution of Nif+ with WT *nifL* and NifA_E356K | This study |
| Av_698 | Derived from MB2302, selected for reconstitution of Nif+ with WT *nifL* and NifA_E366K | This study |

**Table S2.** Plasmids used in this study.

| Plasmid | Description | Reference |
|---|---|---|
| pPR34 | $Cb^R$. *A. vinelandii nifLA* translated from the natural ribosome binding site of *nifL* in pT7-7. | (5) |
| pPMA | $Cb^R$. Derived from pPR34, but encoding *nifA-E356K* | (6) |
| pRT22 | $Cm^R$. *pnifH::lacZ* in pACYC184 | (7) |
| pK18mobsacB | $Km^R$, Mob. Suicide vector for gene replacement. *sacB* gene for counter selection | (8) |
| pMB1816 | $Cb^R$, $Tp^R$. *A. vinelandii* P*nifH::lacZ* fusion flanked by homology regions for integration into *algU* genome locus | (9) |
| pGLR2 | $Km^R$, oriRK2, oriT. pSEVA221-derivative with dual *gfp-luxCDABE* reporter system | (10) |
| pMB2220 | SmR. Strep-Av-NifL (Genscript synthesized, see file S1) cloned into pCDFDuet cut with NcoI/PstI | This study |
| pMB2222 | SmR. Av-SUMO-NifA-6His (Genscript synthesized, see file S1) cloned into pMB2220 cut with NdeI/BglII | This study |
| pMB2223 | CbR. NcoI/PstI fragment from pMB2220 into pETDuet cut with same enzymes | This study |
| pMB2224 | CbR. NdeI/BglII fragment from pMB2222 into pETDuet cut with same enzymes | This study |
| pMB2225 | CbR. NdeI/BglII fragment from pMB2224 into pMB2223 cut with same enzymes | This study |
| pMB2225-S | CbR. PCR generated Av-NifA6his cloned into pMB2223 cut NdeI/BglII | This study |
| pMB2234 | CbR. Av-Strep-NifL269-519 (Twist synthesized, see file S1) cloned by Hi-Fi assembly into pETDuet cut with NcoI/PstI | This study |
| pMB2236 | CbR. Av-NifA188-445-6His (Twist synthesized, see file S1) cloned by Hi-Fi assembly into pMB2234 cut with NdeI/BglII | This study |
| pMB2239 | KmR. Av-NifA188-445-6His amplified by PCR to introduce NcoI/PstI sites cloned into pRSFDuet cut with the same enzymes. | This study |
| pMB2241 | CmR. Av-Strep-NifL269-519 cloned into pACYCDuet cut with NcoI/PstI. | This study |
| pMB23.224 | $Cb^R$. Derived from pPR34, but encoding *nifL-E298A* | This study |
| pMB23.18 | $Cb^R$. Derived from pPR34, but encoding *nifL-E298K* | This study |
| pMB23.408 | $Cb^R$. Derived from pPR34, but encoding *nifL-Q308A* | This study |
| pMB23.416 | $Cb^R$. Derived from pPR34, but encoding *nifL-G309A* | This study |

| | | |
|---|---|---|
| pMB23.93 | $Cb^R$. Derived from pPR34, but encoding *nifL-N312A* | This study |
| pMB23.64 | $Cb^R$. Derived from pPR34, but encoding *nifL-R319A* | This study |
| pMB23.159 | $Cb^R$. Derived from pPR34, but encoding *nifL-R319E* | This study |
| pMB23.104 | $Cb^R$. Derived from pPR34, but encoding *nifL-R323A* | This study |
| pMB23.163 | $Cb^R$. Derived from pPR34, but encoding *nifL-R323E* | This study |
| pMB23.116 | $Cb^R$. Derived from pPR34, but encoding *nifL-R324A* | This study |
| pMB23.520 | $Cb^R$. Derived from pPR34, but encoding *nifL-R324E* | This study |
| pMB23.424 | $Cb^R$. Derived from pPR34, but encoding *nifL-F465A* | This study |
| pMB23.20 | $Cb^R$. Derived from pPR34, but encoding *nifL-E466A* | This study |
| pMB23.393 | $Cb^R$. Derived from pPR34, but encoding *nifL-E466K* | This study |
| pMB23.432 | $Cb^R$. Derived from pPR34, but encoding *nifL-P467A* | This study |
| pMB23.200 | $Cb^R$. Derived from pPR34, but encoding *nifL-Q487A* | This study |
| pMB23.85 | $Cb^R$. Derived from pPR34, but encoding *nifL-E488A* | This study |
| pMB23.400 | $Cb^R$. Derived from pPR34, but encoding *nifL-E488K* | This study |
| pMB23.302 | $Cb^R$. Derived from pPR34, but encoding *nifA-Q224A* | This study |
| pMB23.308 | $Cb^R$. Derived from pPR34, but encoding *nifA-R226D* | This study |
| pMB23.169 | $Cb^R$. Derived from pPR34, but encoding *nifA-K230A* | This study |
| pMB23.178 | $Cb^R$. Derived from pPR34, but encoding *nifA-K230E* | This study |
| pMB23.316 | $Cb^R$. Derived from pPR34, but encoding *nifA-W231A* | This study |
| pMB23.326 | $Cb^R$. Derived from pPR34, but encoding *nifA-N232A* | This study |
| pMB23.330 | $Cb^R$. Derived from pPR34, but encoding *nifA-P257A* | This study |

| | | |
|---|---|---|
| pMB23.490 | $Cb^R$. Derived from pPR34, but encoding *nifA-R258A* | This study |
| pMB23.338 | $Cb^R$. Derived from pPR34, but encoding *nifA-R258D* | This study |
| pMB23.500 | $Cb^R$. Derived from pPR34, but encoding *nifA-E356A* | This study |
| pMB23.504 | $Cb^R$. Derived from pPR34, but encoding *nifA-E360A* | This study |
| pMB23.441 | $Cb^R$. Derived from pPR34, but encoding *nifA-E360K* | This study |
| pMB23.511 | $Cb^R$. Derived from pPR34, but encoding *nifA-E366A* | This study |
| pMB23.350 | $Cb^R$. Derived from pPR34, but encoding *nifA-E366K* | This study |
| pMB23.358 | $Cb^R$. Derived from pPR34, but encoding *nifA-Y369A* | This study |
| pMB23.448 | $Cb^R$. Derived from pPR34, but encoding *nifA-Y369F* | This study |
| pMB23.364 | $Cb^R$. Derived from pPR34, but encoding *nifA-Y370A* | This study |
| pMB23.456 | $Cb^R$. Derived from pPR34, but encoding *nifA-Y370F* | This study |
| pMB23.371 | $Cb^R$. Derived from pPR34, but encoding *nifA-N373A* | This study |
| pMB23.381 | $Cb^R$. Derived from pPR34, but encoding *nifA-V374A* | This study |
| pMB23.385 | $Cb^R$. Derived from pPR34, but encoding *nifA-M375A* | This study |
| pMB23.524 | $Cb^R$. Derived from pPR34 but $NifL_{454-519}$ (ΔNifL) | This study |
| pMB23.590 | $Cb^R$. Derived from pMB23.524, but encoding *nifA-P257A* | This study |
| pMB23.607 | $Cb^R$. Derived from pMB23.524, but encoding *nifA-R258A* | This study |
| pMB23.595 | $Cb^R$. Derived from pMB23.524, but encoding *nifA-Y369A* | This study |
| pMB23.577 | $Cb^R$. Derived from pMB23.524, but encoding *nifA-Y370A* | This study |
| pMB23.581 | $Cb^R$. Derived from pMB23.524, but encoding *nifA-N373A* | This study |
| pMB23.593 | $Cb^R$. Derived from pMB23.524, but encoding *nifA-V374A* | This study |

| | | |
|---|---|---|
| pMB23.598 | $Cb^R$. Derived from pMB23.524, but encoding *nifA-M375A* | This study |
| pKS1901 | $Cb^R$, $Tp^R$. Derived from pMB1816, but promoterless *gfp-luxCDABE* translational fusion derived from pGLR2 | This study |
| pKS1909 | $Cb^R$, $Tp^R$. Derived from pKS1901, but *PnifH* cloned into EcoRI/SalI sites | This study |
| pMB2245 | $Km^R$. nifLA deletion fragment into pK18mobsacBKm | This study |
| pMB23.646 | $Km^R$. XbaI fragment from pPR34 (WT *nifL, nifA*) into pK18mobsacBKm | This study |
| pMB23.614 | $Km^R$. XbaI fragment from pPMA (WT *nifL*, *nifA-E356K*) into pK18mobsacBKm | This study |
| pMB23.673 | $Km^R$. XbaI fragment from pMB23.159 (*nifL-R319E*, WT *nifA*) into pK18mobsacBKm | This study |
| pMB23.622 | $Km^R$. XbaI fragment from pMB23.163 (*nifL-R323E*, WT *nifA*) into pK18mobsacBKm | This study |
| pMB23.716 | KmR. XbaI fragment from pMB23.20 (*nifL-E466A*, WT *nifA*) into pK18mobsacBKm | This study |
| pMB23.658 | $Km^R$. XbaI fragment from pMB23.393 (*nifL-E466K,* WT *nifA*) into pK18mobsacBKm | This study |
| pMB23.632 | $Km^R$. XbaI fragment from pMB23.400 (*nifL-E488K*, WT *nifA*) into pK18mobsacBKm | This study |
| pMB23.663 | $Km^R$. XbaI fragment from pMB23.424 (*nifL-F465A*, WT *nifA*) into pK18mobsacBKm | This study |
| pMB23.686 | $Km^R$. XbaI fragment from pMB23.178 (WT *nifL*, *nifA-K230E*) into pK18mobsacBKm | This study |
| pMB23.718 | $Km^R$. XbaI fragment from pMB23.338 (WT *nifL*, *nifA-R258D*) into pK18mobsacBKm | This study |
| pMB23.698 | $Km^R$. XbaI fragment from pMB23.350 (WT *nifL*, *nifA-E366K*) into pK18mobsacBKm | This study |

**Table S3.** Amino acid sequences used for AlphaFold2 modeling.

| Organism | NifL | | | NifA | | |
|---|---|---|---|---|---|---|
| | aa seq | start | end | aa seq | start | end |
| *Azotobacter vinelandii* DJ | MTPANPTLSNEPQAPHAESDELLP<br>EIFRQTVEHAPIAISITDLKANIL<br>YANRAFRTITGYGSEEVLGKNESI<br>LSNGTTPRLVYQALWGRLAQKKPW<br>SGVLVNRRKDKTLYLAELTVAPVL<br>NEAGETIYYLGMHRDTSELHELEQ<br>RVNNQRLMIEAVVNAAPAAMVVLD<br>RQHRVMLSNPSFCRLARDLVEDGS<br>SESLVALLRENLAAPFETLENQGS<br>AFSGKEISFDLGGRSPRWLSCHGR<br>AIHIENEQAHVFFAPTEERYLLLT<br>INDISELRQKQQDSRLNALKALMA<br>EEELLEGMRETFNAAIHRLQGPVN<br>LISAAMRMLERRLGDKAGNDPVLS<br>AMREASTAGMEALENLSGSIPVRM<br>AESKMPVNLNQLIREVITLCTDQL<br>LAQGIVVDWQPALRLPWVMGGESS<br>LRSMIKHLVDNAIESMSQNQVSRR<br>ELFISTRVENHLVRMEITDSGPGI<br>PPDLVLKVFEPFFSTKPPHRVGRG<br>MGLPVVQEIVAKHAGMVHVDTDYR<br>EGCRIVVELPFSAST | 1 | 519 | MNATIPQRSAKQNPVELYDLQLQA<br>LASIARTLSREQQIDELLEQVLAV<br>LHNDLGLLHGLVTISDPEHGALQI<br>GAIHTDSEAVAQACEGVRYRSGEG<br>VIGNVLKHGNSVVLGRISADPRFL<br>DRLALYDLEMPFIAVPIKNPEGNT<br>IGVLAAQPDCRADEHMPARTRLLE<br>IVANLLAQTVRLVVNIEDGREAAD<br>ERDELRREVRGKYGFENMVVGHTP<br>TMRRVFDQIRRVAKWNSTVLVLGE<br>SGTGKELIASAIHYNSPRAHRPFV<br>RLNCAALPETLLESELFGHEKGAF<br>TGAVKQRKGRFEQADGGTLFLDEI<br>GEISPMFQAKLLRVLQEGEFERVG<br>GNQTVRVNVRIVAATNRDLESEVE<br>KGKFREDLYYRLNVMAIRIPPLRE<br>RTADIPELAEFLLGKIGRQQGRPL<br>TVTDSAIRLLMSHRWPGNVRELEN<br>CLERSAIMSEDGTITRDVVSLTGV<br>DNESPPLAAPLPEVNLADETLDDR<br>ERVIAALEQAGWVQAKAARLLGMT<br>PRQIAYRIQTLNIHMRKI | 1 | 522 |
| *Azotobacter vinelandii* DJ | SELRQKQQDSRLNALKALMAEEEL<br>LEGMRETFNAAIHRLQGPVNLISA<br>AMRMLERRLGDKAGNDPVLSAMRE<br>ASTAGMEALENLSGSIPVRMAESK<br>MPVNLNQLIREVITLCTDQLLAQG<br>IVVDWQPALRLPWVMGGESSLRSM<br>IKHLVDNAIESMSQNQVSRRELFI<br>STRVENHLVRMEITDSGPGIPPDL<br>VLKVFEPFFSTKPPHRVGRGMGLP<br>VVQEIVAKHAGMVHVDTDYREGCR<br>IVVELPFSAST | 269 | 519 | REAADERDELRREVRGKYGFENMV<br>VGHTPTMRRVFDQIRRVAKWNSTV<br>LVLGESGTGKELIASAIHYNSPRA<br>HRPFVRLNCAALPETLLESELFGH<br>EKGAFTGAVKQRKGRFEQADGGTL<br>FLDEIGEISPMFQAKLLRVLQEGE<br>FERVGGNQTVRVNVRIVAATNRDL<br>ESEVEKGKFREDLYYRLNVMAIRI<br>PPLRERTADIP | 188 | 390 |
| *Azoarcus olearius* DQS4 | SASKRQQEALRLAAMRAMLSEGEL<br>NQSLREALAGAVFQLQGPVNLISA<br>AAGLQRRRAGPGAGSDPLARALSE<br>AQRAGEEAIATLQAAMPPEPEEPS<br>GPVNLNEVLRDVLMLETDALLSAG<br>ITVDWHPAHVLPTVQGDPTALRTL<br>FRQLVSNAIEAMNVRGWKERALVL<br>RTSTQNGQVDVEVTDTGPGIPEAL<br>RLKVFEPFFSTKKARAAGRGVGLS<br>LAQEIVTRHRGMLEIDPEYSGGCR<br>LRISFPALRSAPLDLEGGR | 260 | 518 | RDIAEERDNLRRTVRNRFGFDNIV<br>GHTQRMRQVFEQIRQVAKWNTTVL<br>VRGETGTGKELIAQAIHYNSPRAA<br>GPFVKLNCAALPENLLESELFGHE<br>KGAFTGALTQRKGRFEMADGGTLF<br>LDEIGEISASFQAKLLRVLQEGEL<br>ERVGGARTLRVDVRVIAATNRDLE<br>LEVEAGKFREDLYYRLNVMPIMLP<br>PLRERVEDIP | 184 | 385 |
| *Azonexus hydrophilus* | TNLREEQERAKGAALKASLAEEER<br>TAAIREGLSAAVFRLEEPLNVMAS<br>AVAVLQRRDPASAEMLRHALDNSR<br>EHIESLRQVIPQSPHEILIRVNLN<br>EILRDVLEVCTPRLLSAGITVDWQ<br>PSAVLPPLNGRPLQLRMLFKALVE<br>NAIEAMNVKGWPRRELQIKSEVVD<br>DCLRVAILDSGPGIPMEWRHKAFE<br>PFFTAKNGSGKHIGTGLSRAQQVV<br>ADHGGIIDLDESPAGGCAAIVEFR<br>VDGDPI | 261 | 506 | STLLEERDLLRRTVRHQHGFDSMV<br>GRSAVMRRVFEQARMVAKWNTTVL<br>IRGETGTGKELIANAIHYNSPRAR<br>NMLVKLNCAALPENLLESELFGHE<br>RGAFTGAVEARKGRFEQANGGTLF<br>LDEIGEVSAAFQAKLLRILQEGEF<br>ERVGGSKTIKVDVRIIAATHRDLE<br>TAVEMGDFREDLFYRLNVMPLFLP<br>PLRERIEDIP | 193 | 394 |

| | | | | | | |
|---|---|---|---|---|---|---|
| *Ectothiorhodospira marina* DSM241 | TELKRQQEEMRMNALRALTAESEL<br>VESMRETLTGAIYQMQGPVNMIAA<br>ATQLLERRQGEGATDPLLTVLQDA<br>LASGRQALQTLENCVPEERDEAIA<br>PVNLNGLLREVLGLSTRRLLTSGV<br>VVDWDPAPVLPNVLGREGRLRGMF<br>KQLVDNALDALDDSGTGQREIRIA<br>TRALEDGVQVIIQDSGPGIPESMR<br>LRVFEPFVSTKGRSGGGAGMGLPM<br>VQEVVNQHAGLVEVDPSCQQGCRI<br>RLEFPNASQRKL | 271 | 522 | RALTDERDRLRRQVRGGYHLDNVV<br>GHSKEMRRVFEQVRQVAKWHTTVL<br>VRGETGTGKELIANAIHYNSPRAN<br>GHFVKLNCAALPDNLLESELFGHE<br>RGAFTGAVSQRKGRFEQADGGTIF<br>LDEIGEISQAFQAKLLRVLQEGEF<br>ERVGGTRTLKVDVRVIAATNRPLE<br>EDVERGDFREDLYYRLNVMPIFVP<br>ALRNRSEDIP | 187 | 388 |
| *Mariprofundus ferrooxydans* M34 | TGLKAHEHAVRCNTMRTILAKQEV<br>VQSLREILNGAAYQLQVPVNVLSA<br>VTNMLKQREDKGGCNSSMLEALDQ<br>ALDSGRKALATLDSSKPTILTEAP<br>VDVDLNELLRDVVALSEERVSDMG<br>VVIDWQQFEALPTMKGQQWQLRGA<br>LKQLVDNALDAMTQVDGGHREIQL<br>KTELCDDMIKVHICDSGPGIPDEL<br>RRSIFEPFFTTKRGESSAGMGLPT<br>CQSVIREHGGMIWVENSMLGGICM<br>RLQFPLVDAEKQG | 251 | 503 | QALTSERDQLRQAVQVNYGFDNIV<br>GHTDIMRRVFEQVQQVSKWNTTVL<br>IRGESGTGKELIANAIHYNSPRAS<br>DPFIKLNCAALPDNLLESELFGHE<br>KGAFTGAVNQRKGRFEQADGGTIF<br>LDEIGEISGTFQAKLLRVLQEGEF<br>ERVGGTRTMQVDVRVVAATNKNLE<br>QEVEEGGFREDLYYRLNVMPINLP<br>SLRERAEDIP | 187 | 388 |
| *Martelella endophytica* YC6887 | TSLKWQYEQVRLNAVRAQMAELAL<br>RRGTNEIIEGALYQLQGPLNVMQA<br>MSGMVKRREGEDSHMGMAIDEALS<br>SGNRAIDRLKLSQPPLPQLAPAPI<br>NLNQVLRDVLVLSANRLSASGVVV<br>DWRPERELPSIYGHENLLRILMKL<br>LLDNAITAVGEPGAYAREILIATR<br>IEGPSMIEIFVRDRGPGVDKAIRS<br>QIFEPFFSAWSPKRRGAGMGLAIG<br>QQIVSELGGEMMIETAHQPGFGMC<br>VALPTCGALKNGT | 294 | 546 | DLLRTERDQLRRHVKQQHGFDNII<br>GATESMRVVFDQVRVVAKWNTTVL<br>IRGESGTGKELIANAIHYNSPRSS<br>AQFLKLNCAALPDNLLESELFGHE<br>KGAFTGATAQRKGRFEQADGGTLF<br>LDEIGEISPAFQSKLLRVLQEGEF<br>ERVGGNRTLKVDVRIIAATNKDLE<br>TAVENGDFREDLYYRLNVMALTMP<br>ALRERIDDIP | 195 | 396 |
| *Pseudomonas stutzeri* A1501 | TGLRQQQQASQLNALKVLMAEEEL<br>LDGMRETFNGAIHRLQGPVNLISA<br>ALRMLERRLGDSAEGDPVLSAMRE<br>ASQAGMDALESLSGSIPQRRAGGC<br>VPVNINQLIREVVSLMTDQLLAQG<br>IVVDWQPALRLPWVMGAEGRLRSM<br>IKQLLENAIEAMSQNQDNPRTLSI<br>VTRVQGQRVVRLEIADSGPGIAPE<br>LALKVFEPFFSTKPPHKTGRGMGL<br>AMVQETVTEHAGTVHIDSGYDQGC<br>RIVVELPFSAS | 287 | 537 | QEVVDERDELRREVRAKYGFENMV<br>VGHTASMRRVFDQVRRVAKWNSTV<br>LILGESGTGKELIASAIHYNSPRA<br>HQPLVRLNCAALPETLLESELFGH<br>EKGAFTGAVKQRKGRFEQADGGTL<br>FLDEIGEISPMFQAKLLRVLQEGE<br>LERVGGSQTVKVNVRIVAATNRDL<br>EHEVEQGKFREDLYYRLNVMAIRV<br>PPLRERSADIP | 188 | 390 |
| *Sideroxydans lithotrophicus* ES-1 | SEIKRQQEALRINSLRALLAEQER<br>IQGLRETLAGAVYQLEGPLNMLSA<br>AANMMERRREAGADLPAASALEEA<br>MQKSREVLETLRGCIPEQGNEALQ<br>PVNLNEVLADVLKLTTASLLGAGI<br>VVEWSPSNEPPVVSGHPGQLTNLF<br>KQLISNAVEAVNERRGGQRELRIA<br>CTSRGDHLEVFVEDSGPGIADDLR<br>YKVFQPFFTTKGAARQHLGLGLAM<br>AQEVVIRHGGTIDIDPGYHAGCRV<br>RVQLPLLTGANHG | 269 | 521 | NEIAEERDVLRRTVRGQYGFDNII<br>GHTQPMRRIFEQVRLVAKWNTTVL<br>IRGETGTGKELIANAIHYNSPRAR<br>GSFVKLNCAALPETLLESELFGHE<br>KGAFTGAVAQRKGRFEQAEGGTVF<br>LDEIGEISPSFQAKLLRVLQEGEL<br>ERVGGTKTIKVDVRVVAATHRDLE<br>AEVEKGRFREDLYYRLNVMPLYLP<br>PLRDRMEDIP | 184 | 385 |
| *Klebisiella oxytoca* M5a1 | TQQRQQQEQGRLDRLKQQMTAGKL<br>LAAIRESLDAALIQLNCPINMLAA<br>ARRLNGEGSGNVALDAAWREGEEA<br>MARLQRCRPSLELESNAVWPLQPF<br>FDDLYALYRTRFDDRARLQVDMAS<br>PHLVGFGQRTQLLACLSLWLDRTL<br>ALAAELPSVPLEIELYAEEDEGWL<br>SLYLNDNVPLLQVRYAHSPDALNS<br>PGKGMELRLIQTLVAYHRGAIELA<br>SRPQGGTSLVLRFPLFNTLTGGEQ | 256 | 496 | QSPRIERPRACTPSRGFGLENMVG<br>KSPAMRQIMDIIRQVSRWDTTVLV<br>RGESGTGKELIANAIHHNSPRAAA<br>AFVKFNCAALPDNLLESELFGHEK<br>GAFTGAVRQRKGRFELADGGTLFL<br>DEIGESSASFQAKLLRILQEGEME<br>RVGGDETLRVNVRIIAATNRHLEE<br>EVRLGHFREDLYYRLNVMPIALPP<br>LRERQEDIA | 191 | 391 |

**Table S4.** Summary of cryo-EM data collection, processing, refinement and validation

| | |
|---|---|
| **Data collection and processing** | |
| Microscope | Titan Krios |
| Detector | Gatan K3 Summit |
| Detector mode | counting; super resolution; binning x2 |
| Magnification | 105,000x |
| Magnified/extracted pixel size (Å) | 0.831/1.484 |
| Voltage (kV) | 300 |
| Defocus range ($\mu$m) | -0.7 to -2.2 |
| Movie format | TIFF |
| Movies collected/used | 14,218/13,798 |
| Exposure time per movie (sec) | 2.27 |
| Total dose per movie ($e^-$/Å$^2$) | 50.0 |
| Frames per movie | 50 |
| Particle images | 209,009 |
| Extracted particle box size (pixels) | 168 |
| Symmetry imposed | None (C1) |
| Resolution (global; Å): | |
| FSC 0.5 (unmasked/masked) | 9.30/7.79 |
| FSC 0.143 (unmasked/masked) | 7.74/6.45 |
| Resolution range (local; Å) | 3.98 – 92.40 |
| Resolution range due to anisotropy (Å) | 4.62 – 9.03 |
| **Refinement** | |
| Initial model used | AlphaFold3 |
| Map sharpening *B* factor (Å$^2$) | -458.5 |
| Non-hydrogen atoms | 9,944 |
| Protein residues | 1,243 |
| Residues present | NifL: 21-517 x2; NifA: 210-458 |
| Ligands | FAD x2; ADP x3 |
| **R.m.s. deviations** | |
| Bond lengths (Å) | 0.003 |
| Bond angles (°) | 0.598 |
| **Validation** | |
| Molprobity score | 1.85 |
| Clashscore | 16.3 |
| CaBLAM outliers (%) | 0.81 |
| Rotamer outliers (%) | 0.0 |
| Cβ outliers (%) | 0.0 |
| Ramachandran plot: | |
| Favoured (%) | 97.3 |
| Allowed (%) | 2.7 |
| Disallowed (%) | 0.0 |
| Overall Z-score | 0.78 ± 0.25 |
| Fit to map ($CC_{mask}$) | 0.69 |
| Map to model FSC (Å) | |

| | |
|---|---|
| FSC 0.5 (unmasked/masked) | 9.9/9.7 |
| FSC 0.143 (unmasked/masked) | 6.9/6.7 |
| **Accession codes** | |
| EMPIAR (raw data) | ________ |
| EMDB (maps) | EMD-53294 |
| PDB (model) | 9QQ6 |

**Supplementary File S1**. Nucleotide sequences of synthesized genes used in this study followed by their respective amino acid sequences.

**A-1)** DNA sequence of synthesized Av-Strep-NifL

Underlined: NcoI and PstI restriction sites
Green: Strep-Tag II
Red: enterokinase site
Blue: codon optimized *nifL*

CCATGGCAAGCTGGAGCCACCCGCAGTTCGAAAAGGGTGCAGATGACGACGACAAGGTAACCCCGGCGAAC
CCGACCCTGAGCAACGAACCGCAAGCGCCACATGCGGAAAGCGATGAACTGCTGCCGGAAATTTTTCGTCA
GACCGTGGAACATGCGCCGATTGCGATTAGCATTACCGATCTGAAAGCGAACATTCTGTATGCGAACCGCG
CGTTTCGCACCATTACCGGCTATGGCAGCGAAGAAGTGCTGGGCAAAAACGAAAGCATTCTGAGCAACGGC
ACCACCCCGCGCCTGGTGTATCAAGCGCTGTGGGGGCGCTTAGCGCAAAAGAAACCGTGGAGCGGCGTGCT
GGTGAACCGCCGCAAAGATAAAACCCTGTATCTGGCGGAACTGACCGTGGCGCCGGTGCTGAACGAAGCGG
GCGAAACCATTTATTATCTGGGTATGCATCGCGATACGAGCGAACTGCATGAACTGGAACAGCGCGTGAAC
AATCAGCGCCTGATGATTGAAGCGGTGGTGAACGCGGCGCCGGCGGCGATGGTGGTGCTGGATCGTCAGCA
TCGCGTGATGCTGAGCAACCCGAGCTTTTGCCGCCTGGCGCGAGACCTGGTGGAAGATGGCAGCAGCGAAA
GCCTGGTGGCGCTGCTGCGCGAAAACCTGGCGGCGCCGTTTGAAACCCTGGAAAACCAAGGCAGCGCGTTT
AGCGGCAAAGAAATTAGCTTTGATCTGGGTGGCCGCAGCCCGCGCTGGCTGAGCTGTCATGGCCGCGCGAT
TCATATTGAAAACGAACAAGCGCATGTGTTTTTTGCGCCGACCGAAGAACGCTATCTGCTGCTGACCATTA
ACGATATTAGCGAACTGCGTCAGAAACAGCAAGATAGCCGCCTGAACGCGCTGAAAGCGCTGATGGCGGAA
GAAGAACTGCTGGAAGGTATGCGCGAAACCTTTAACGCGGCGATTCATCGCCTGCAAGGCCCCGTGAACCT
GATTAGCGCGGCGATGCGTATGCTGGAACGCCGCCTGGGCGATAAAGCGGGCAACGATCCGGTGCTGAGCG
CGATGCGCGAAGCGAGCACGGCGGGCATGGAAGCGCTGGAAAACCTGAGCGGCAGCATTCCGGTGCGCATG
GCGGAAAGCAAAATGCCGGTGAACCTGAATCAGCTGATTCGCGAAGTGATTACCCTGTGCACCGATCAGCT
GCTGGCGCAAGGCATAGTAGTCGATTGGCAGCCGGCGCTGCGCCTCCCGTGGGTGATGGGCGGCGAAAGCA
GCCTGCGCAGCATGATTAAACATCTGGTGGATAACGCGATTGAAAGCATGAGTCAGAACCAAGTGAGCCGC
CGCGAACTGTTTATTAGCACCCGCGTGGAAAACCATCTGGTGCGCATGGAAATTACCGATAGCGGTCCGGG
CATTCCGCCGGATCTGGTGCTGAAAGTGTTTGAACCGTTTTTTAGCACCAAACCGCCGCATCGCGTGGGTC
GCGGCATGGGCCTGCCGGTGGTGCAAGAAATTGTGGCGAAACATGCGGGCATGGTGCATGTGGATACCGAT
TATCGCGAAGGCTGCCGCATTGTGGTGGAACTGCCGTTTAGCGCGAGCACCTAACTGCAG

**A-2)** Amino acid sequence of Av-Strep-NifL

Green: Strep-Tag II
Red: enterokinase site
Blue: Av-NifL

MASWSHPQFEKGADDDDKVTPANPTLSNEPQAPHAESDELLPEIFRQTVEHAPIAISITDLKANILYANRA
FRTITGYGSEEVLGKNESILSNGTTPRLVYQALWGRLAQKKPWSGVLVNRRKDKTLYLAELTVAPVLNEAG
ETIYYLGMHRDTSELHELEQRVNNQRLMIEAVVNAAPAAMVVLDRQHRVMLSNPSFCRLARDLVEDGSSES
LVALLRENLAAPFETLENQGSAFSGKEISFDLGGRSPRWLSCHGRAIHIENEQAHVFFAPTEERYLLLTIN
DISELRQKQQDSRLNALKALMAEEELLEGMRETFNAAIHRLQGPVNLISAAMRMLERRLGDKAGNDPVLSA
MREASTAGMEALENLSGSIPVRMAESKMPVNLNQLIREVITLCTDQLLAQGIVVDWQPALRLPWVMGGESS
LRSMIKHLVDNAIESMSQNQVSRRELFISTRVENHLVRMEITDSGPGIPPDLVLKVFEPFFSTKPPHRVGR
GMGLPVVQEIVAKHAGMVHVDTDYREGCRIVVELPFSAST*

**B-1)** DNA sequence of synthesized Av-SUMO-NifA-6xHis
Underlined: NdeI and BglII restriction sites

Green: SUMO
Blue: codon optimized *nifA*
Orange: 6xHis

```
CATATGTCGGACTCAGAAGTCAATCAAGAAGCTAAGCCAGAGGTCAAGCCAGAAGTCAAGCCTGAGACTCA
CATCAATTTAAAGGTGTCCGATGGATCTTCAGAAATCTTCTTCAAGATCAAAAAGACCACTCCTTTAAGAA
GGCTGATGGAAGCGTTCGCTAAAAGACAGGGTAAGGAAATGGACTCCTTAAGATTCTTGTACGACGGTATT
AGAATCCAAGCTGATCAGACCCCTGAAGATTTGGACATGGAGGATAACGATATTATTGAGGCTCACAGAGA
ACAGATTGGTGGTAACGCGACCATTCCGCAGCGCAGCGCGAAACAGAACCCGGTGGAATTATATGATTTAC
AGCTGCAAGCGCTGGCGAGCATTGCGCGCACCCTGAGCCGCGAACAGCAGATTGATGAACTGCTGGAACAA
GTGCTGGCGGTGCTGCATAACGATCTGGGCCTGCTGCATGGCCTGGTGACCATTAGCGATCCGGAACATGG
CGCGTTACAGATTGGCGCGATTCATACCGATAGCGAAGCGGTGGCGCAAGCGTGCGAAGGCGTGCGCTATC
GCAGCGGCGAAGGCGTGATTGGCAACGTGCTGAAACATGGCAACAGCGTGGTGCTGGGCCGCATTAGCGCC
GATCCGCGCTTTCTGGATCGCCTGGCGCTCTATGATCTGGAAATGCCGTTTATTGCGGTGCCGATTAAAAA
CCCGGAAGGCAACACCATTGGCGTGCTGGCGGCGCAGCCGGATTGCCGCGCGGATGAACACATGCCGGCGC
GCACCCGCCTGCTGGAAATTGTGGCGAACCTGCTGGCGCAGACAGTGCGCCTGGTGGTGAACATTGAAGAT
GGCCGCGAAGCGGCGGATGAACGCGATGAACTGCGCCGCGAAGTGCGCGGCAAATATGGCTTTGAAAACAT
GGTGGTGGGCCATACCCCGACCATGCGCCGCGTGTTTGATCAGATTCGCCGCGTGGCGAAATGGAACAGCA
CCGTGCTGGTGCTGGGGGAAAGTGGCACCGGCAAAGAACTGATTGCGAGCGCGATTCACTATAACAGCCCG
CGCGCGCATCGCCCGTTTGTGCGCCTGAACTGCGCGGCGCTGCCGGAAACCCTGCTTGAAAGCGAACTTTT
TGGCCATGAAAAAGGCGCGTTTACCGGCGCGGTGAAACAGCGCAAAGGCCGCTTTGAACAAGCGGATGGCG
GCACCCTGTTTCTGGATGAAATTGGCGAAATTAGCCCGATGTTTCAAGCGAAACTGCTGCGCGTGCTGCAA
GAAGGCGAATTTGAACGCGTGGGCGGCAATCAGACCGTGCGCGTGAACGTGCGCATTGTGGCGGCGACCAA
CCGCGATCTAGAGAGCGAAGTGGAAAAAGGCAAATTTCGCGAGGATCTGTATTATCGCCTGAACGTGATGG
CGATTCGCATTCCGCCGCTGCGCGAACGCACGGCGGATATTCCGGAACTGGCGGAATTTCTGCTGGGCAAA
ATTGGCCGTCAGCAAGGCCGCCCGCTGACCGTGACCGATAGCGCGATTCGCCTGCTGATGAGCCATCGCTG
GCCGGGCAACGTGCGCGAACTGGAAAACTGCCTGGAACGCAGCGCGATTATGAGCGAAGATGGCACCATTA
CCCGCGATGTGGTGAGCCTGACCGGCGTGGATAACGAAAGCCCGCCGCTGGCGGCGCCGCTGCCGGAAGTG
AACCTGGCGGATGAAACCCTGGATGATCGCGAACGCGTGATTGCGGCGCTCGAACAAGCGGGATGGGTTCA
AGCGAAGGCGGCTCGCCTGCTGGGCATGACCCCGCGTCAGATTGCGTATCGCATTCAGACCCTGAACATTC
ACATGCGCAAAATTCATCATCATCATCATCACTAAAGATCT
```

**B-2)** Amino acid sequence of Av-SUMO-NifA-6xHis

Green: SUMO
Blue: Av-NifA
Orange: 6xHis

```
MSDSEVNQEAKPEVKPEVKPETHINLKVSDGSSEIFFKIKKTTPLRRLMEAFAKRQGKEMDSLRFLYDGIR
IQADQTPEDLDMEDNDIIEAHREQIGGNATIPQRSAKQNPVELYDLQLQALASIARTLSREQQIDELLEQV
LAVLHNDLGLLHGLVTISDPEHGALQIGAIHTDSEAVAQACEGVRYRSGEGVIGNVLKHGNSVVLGRISAD
PRFLDRLALYDLEMPFIAVPIKNPEGNTIGVLAAQPDCRADEHMPARTRLLEIVANLLAQTVRLVVNIEDG
REAADERDELRREVRGKYGFENMVVGHTPTMRRVFDQIRRVAKWNSTVLVLGESGTGKELIASAIHYNSPR
AHRPFVRLNCAALPETLLESELFGHEKGAFTGAVKQRKGRFEQADGGTLFLDEIGEISPMFQAKLLRVLQE
GEFERVGGNQTVRVNVRIVAATNRDLESEVEKGKFREDLYYRLNVMAIRIPPLRERTADIPELAEFLLGKI
GRQQGRPLTVTDSAIRLLMSHRWPGNVRELENCLERSAIMSEDGTITRDVVSLTGVDNESPPLAAPLPEVN
LADETLDDRERVIAALEQAGWVQAKAARLLGMTPRQIAYRIQTLNIHMRKIHHHHHH*
```

**C-1)** DNA sequence of synthesized Av-NifA-6xHis
Underlined: NdeI and BglII restriction sites

Blue: codon optimized *nifA*
Orange: 6xHis

CATATGAACGCGACCATTCCGCAGCGCAGCGCGAAACAGAACCCGGTGGAATTATATGATTTACAGCTGCA
AGCGCTGGCGAGCATTGCGCGCACCCTGAGCCGCGAACAGCAGATTGATGAACTGCTGGAACAAGTGCTGG
CGGTGCTGCATAACGATCTGGGCCTGCTGCATGGCCTGGTGACCATTAGCGATCCGGAACATGGCGCGTTA
CAGATTGGCGCGATTCATACCGATAGCGAAGCGGTGGCGCAAGCGTGCGAAGGCGTGCGCTATCGCAGCGG
CGAAGGCGTGATTGGCAACGTGCTGAAACATGGCAACAGCGTGGTGCTGGGCCGCATTAGCGCCGATCCGC
GCTTTCTGGATCGCCTGGCGCTCTATGATCTGGAAATGCCGTTTATTGCGGTGCCGATTAAAAACCCGGAA
GGCAACACCATTGGCGTGCTGGCGGCGCAGCCGGATTGCCGCGCGGATGAACACATGCCGGCGCGCACCCG
CCTGCTGGAAATTGTGGCGAACCTGCTGGCGCAGACAGTGCGCCTGGTGGTGAACATTGAAGATGGCCGCG
AAGCGGCGGATGAACGCGATGAACTGCGCCGCGAAGTGCGCGGCAAATATGGCTTTGAAAACATGGTGGTG
GGCCATACCCCGACCATGCGCCGCGTGTTTGATCAGATTCGCCGCGTGGCGAAATGGAACAGCACCGTGCT
GGTGCTGGGGGAAAGTGGCACCGGCAAAGAACTGATTGCGAGCGCGATTCACTATAACAGCCCGCGCGCGC
ATCGCCCGTTTGTGCGCCTGAACTGCGCGGCGCTGCCGGAAACCCTGCTTGAAAGCGAACTTTTTGGCCAT
GAAAAAGGCGCGTTTACCGGCGCGGTGAAACAGCGCAAAGGCCGCTTTGAACAAGCGGATGGCGGCACCCT
GTTTCTGGATGAAATTGGCGAAATTAGCCCGATGTTTCAAGCGAAACTGCTGCGCGTGCTGCAAGAAGGCG
AATTTGAACGCGTGGGCGGCAATCAGACCGTGCGCGTGAACGTGCGCATTGTGGCGGCGACCAACCGCGAT
CTAGAGAGCGAAGTGGAAAAAGGCAAATTTCGCGAGGATCTGTATTATCGCCTGAACGTGATGGCGATTCG
CATTCCGCCGCTGCGCGAACGCACGGCGGATATTCCGGAACTGGCGGAATTTCTGCTGGGCAAAATTGGCC
GTCAGCAAGGCCGCCCGCTGACCGTGACCGATAGCGCGATTCGCCTGCTGATGAGCCATCGCTGGCCGGGC
AACGTGCGCGAACTGGAAAACTGCCTGGAACGCAGCGCGATTATGAGCGAAGATGGCACCATTACCCGCGA
TGTGGTGAGCCTGACCGGCGTGGATAACGAAAGCCCGCCGCTGGCGGCGCCGCTGCCGGAAGTGAACCTGG
CGGATGAAACCCTGGATGATCGCGAACGCGTGATTGCGGCGCTCGAACAAGCGGGATGGGTTCAAGCGAAG
GCGGCTCGCCTGCTGGGCATGACCCCGCGTCAGATTGCGTATCGCATTCAGACCCTGAACATTCACATGCG
CAAAATTCATCATCATCATCATCACTAAAGATCT

**C-2)** Amino acid sequence of Av-NifA-6xHis

Blue: Av-NifA
Orange: 6xHis

MNATIPQRSAKQNPVELYDLQLQALASIARTLSREQQIDELLEQVLAVLHNDLGLLHGLVTISDPEHGALQ
IGAIHTDSEAVAQACEGVRYRSGEGVIGNVLKHGNSVVLGRISADPRFLDRLALYDLEMPFIAVPIKNPEG
NTIGVLAAQPDCRADEHMPARTRLLEIVANLLAQTVRLVVNIEDGREAADERDELRREVRGKYGFENMVVG
HTPTMRRVFDQIRRVAKWNSTVLVLGESGTGKELIASAIHYNSPRAHRPFVRLNCAALPETLLESELFGHE
KGAFTGAVKQRKGRFEQADGGTLFLDEIGEISPMFQAKLLRVLQEGEFERVGGNQTVRVNVRIVAATNRDL
ESEVEKGKFREDLYYRLNVMAIRIPPLRERTADIPELAEFLLGKIGRQQGRPLTVTDSAIRLLMSHRWPGN
VRELENCLERSAIMSEDGTITRDVVSLTGVDNESPPLAAPLPEVNLADETLDDRERVIAALEQAGWVQAKA
ARLLGMTPRQIAYRIQTLNIHMRKIHHHHHH*

**D-1)** DNA sequence of synthesized Av-Strep-$NifL_{269\text{-}519}$

Underlined: NcoI and PstI restriction sites
Green: Strep-Tag II
Red: enterokinase site
Blue: codon optimized *nifL*$_{269\text{-}519}$

CCATGGCAAGCTGGAGCCACCCGCAGTTCGAAAAGGGTGCAGATGACGACGACAAGGTAAGCGAACTGCGT
CAGAAACAGCAAGATAGCCGCCTGAACGCGCTGAAAGCGCTGATGGCGGAAGAAGAACTGCTGGAAGGTAT
GCGCGAAACCTTTAACGCGGCGATTCATCGCCTGCAAGGCCCCGTGAACCTGATTAGCGCGGCGATGCGTA
TGCTGGAACGCCGCCTGGGCGATAAAGCGGGCAACGATCCGGTGCTGAGCGCGATGCGCGAAGCGAGCACG
GCGGGCATGGAAGCGCTGGAAAACCTGAGCGGCAGCATTCCGGTGCGCATGGCGGAAAGCAAAATGCCGGT
GAACCTGAATCAGCTGATTCGCGAAGTGATTACCCTGTGCACCGATCAGCTGCTGGCGCAAGGCATAGTAG
TCGATTGGCAGCCGGCGCTGCGCCTCCCGTGGGTGATGGGCGGCGAAAGCAGCCTGCGCAGCATGATTAAA
CATCTGGTGGATAACGCGATTGAAAGCATGAGTCAGAACCAAGTGAGCCGCCGCGAACTGTTTATTAGCAC
CCGCGTGGAAAACCATCTGGTGCGCATGGAAATTACCGATAGCGGTCCGGGCATTCCGCCGGATCTGGTGC
TGAAAGTGTTTGAACCGTTTTTTAGCACCAAACCGCCGCATCGCGTGGGTCGCGGCATGGGCCTGCCGGTG
GTGCAAGAAATTGTGGCGAAACATGCGGGCATGGTGCATGTGGATACCGATTATCGCGAAGGCTGCCGCAT
TGTGGTGGAACTGCCGTTTAGCGCGAGCACCTAACTGCAG

**D-2)** Amino acid sequence of Av-Strep-$NifL_{269\text{-}519}$

Green: Strep-Tag II
Red: enterokinase site
Blue: Av-$NifL_{269\text{-}519}$

MASWSHPQFEKGADDDDKVSELRQKQQDSRLNALKALMAEEELLEGMRETFNAAIHRLQGPVNLISAAMRM
LERRLGDKAGNDPVLSAMREASTAGMEALENLSGSIPVRMAESKMPVNLNQLIREVITLCTDQLLAQGIVV
DWQPALRLPWVMGGESSLRSMIKHLVDNAIESMSQNQVSRRELFISTRVENHLVRMEITDSGPGIPPDLVL
KVFEPFFSTKPPHRVGRGMGLPVVQEIVAKHAGMVHVDTDYREGCRIVVELPFSAST*

**E-1)** DNA sequence of synthesized Av-NifA$_{188\text{-}445}$-6xHis
Underlined: NdeI and BglII restriction sites

Blue: codon optimized *nifA*$_{188\text{-}445}$
Orange: 6xHis

ATGCGCGAAGCGGCGGATGAACGCGATGAACTGCGCCGCGAAGTGCGCGGCAAATATGGCTTTGAAAACAT
GGTGGTGGGCCATACCCCGACCATGCGCCGCGTGTTTGATCAGATTCGCCGCGTGGCGAAATGGAACAGCA
CCGTGCTGGTGCTGGGGGAAAGTGGCACCGGCAAAGAACTGATTGCGAGCGCGATTCACTATAACAGCCCG
CGCGCGCATCGCCCGTTTGTGCGCCTGAACTGCGCGGCGCTGCCGGAAACCCTGCTTGAAAGCGAACTTTT
TGGCCATGAAAAAGGCGCGTTTACCGGCGCGGTGAAACAGCGCAAAGGCCGCTTTGAACAAGCGGATGGCG
GCACCCTGTTTCTGGATGAAATTGGCGAAATTAGCCCGATGTTTCAAGCGAAACTGCTGCGCGTGCTGCAA
GAAGGCGAATTTGAACGCGTGGGCGGCAATCAGACCGTGCGCGTGAACGTGCGCATTGTGGCGGCGACCAA
CCGCGATCTAGAGAGCGAAGTGGAAAAAGGCAAATTTCGCGAGGATCTGTATTATCGCCTGAACGTGATGG
CGATTCGCATTCCGCCGCTGCGCGAACGCACGGCGGATATTCCGGAACTGGCGGAATTTCTGCTGGGCAAA
ATTGGCCGTCAGCAAGGCCGCCCGCTGACCGTGACCGATAGCGCGATTCGCCTGCTGATGAGCCATCGCTG
GCCGGGCAACGTGCGCGAACTGGAAAACTGCCTGGAACGCAGCGCGATTATGAGCGAAGATGGCACCGGTG
GTCATCATCATCATCATCACTAA

**E-2)** Amino acid sequence of Av-NifA-6xHis

Blue: Av-NifA$_{188\text{-}445}$
Orange: 6xHis

MREAADERDELRREVRGKYGFENMVVGHTPTMRRVFDQIRRVAKWNSTVLVLGESGTGKELIASAIHYNSP
RAHRPFVRLNCAALPETLLESELFGHEKGAFTGAVKQRKGRFEQADGGTLFLDEIGEISPMFQAKLLRVLQ
EGEFERVGGNQTVRVNVRIVAATNRDLESEVEKGKFREDLYYRLNVMAIRIPPLRERTADIPELAEFLLGK
IGRQQGRPLTVTDSAIRLLMSHRWPGNVRELENCLERSAIMSEDGTGGHHHHHH*
